# Organization of mouse prefrontal cortex subnetwork revealed by spatial single-cell multi-omic analysis of SPIDER-Seq

**DOI:** 10.64898/2025.12.29.696799

**Authors:** Leqiang Sun, Hu Zheng, Yayu Huang, Xuehuan Huang, Keji Yan, Zhongchao Wang, Liyao Yang, Yiping Yue, Xiaojuan Gou, Guohua Du, Yang Wang, Xiaofeng Wu, Huazhen Liu, Hang Chen, Daqing Ma, Yunyun Han, Jinxia Dai, Gang Cao

## Abstract

Deciphering the connectome, anatomy, transcriptome and spatial-omics integrated multi-modal brain atlas and the underlying organization principles remains a great challenge. We developed a Single-cell Projectome-transcriptome In situ Deciphering Sequencing (SPIDER-Seq) technique by combining viral barcoding tracing with single-cell sequencing and spatial-omics. This empowers us to delineate a integrated single-cell spatial molecular, cellular, anatomic and projectomic atlas of mouse prefrontal cortex (PFC). The projectomic and transcriptomic cell clusters display distinct modular organization principles, but are coordinately configured in the PFC. The projection neurons gradiently occupied different territories in the PFC aligning with their wiring patterns. Importantly, they show higher co-projection probability to the downstream nuclei with reciprocal circuit connections. Moreover, we integrated projectomic atlas with their distinct spectrum of neurotransmitter/neuropeptide and the receptors-related gene profiles and depicted PFC neural signal transmission network. By which, we uncovered potential mechanisms underlying the complexity and specificity of neural transmission. Finally, leveraging machine learning, we predicted neuron projections with high accuracy by combining gene profiles and spatial information. As a proof of concept, we used this model to predict projections of fear recall engram neurons. This study facilitates our understanding of brain multi-modal network and neural computation.

## INTRODUCTION

Deciphering the sophisticated neural connectome and the underlying wiring logic is essential for understanding brain computation and advancing artificial intelligence [1, 2]. Although the mesoscopic connectivity in several model organisms have been well characterized [3–7], the detailed wiring information at single neuron resolution remains largely unknown [8, 9], impeding the deep understanding of neural computational logic at level of precision. Distinct signaling molecules and wiring-related genes expressed by individual neurons result in their unique and dynamic capacities for information transmission and plasticity. These gene expression profiles, in combination with the neural projection patterns and connection topological organization, make neural networks dynamic and highly integrated complex systems [10]. The integrative analysis of the multi-modal characteristics of these neural network fundamental units with single neuron resolution is essential for understanding the functions of neural network. Thus, high-throughput neuronal connectivity decoding techniques with the capacity to integrate gene expression and spatial distribution with single cell resolution are highly desired.

In the past decade, middle or high-throughput methods for analyzing neuronal connectivity have been developed, such as MAPseq, BARseq, BRICseq and fMOST, revolutionizing the neural network research paradigm and providing new insights into the topological structures of neural circuits [11–15]. BARseq2, an improved version of BARseq, offers potential path to uncovering the molecular logic underlying neuronal circuits [16]. Because the replication of the barcode-carrying Sindbis virus perturbs neuronal transcription, there is a need for combining low-toxicity circuit-tracing virus with single-cell sequencing to simultaneously explore circuit architecture and intact transcriptomics. Technologies such as VECTORseq, Retro-Seq, and Epi-Retro-Seq have been developed to decipher transcriptomes and even epigenomes in specific circuits [17–20]. However, these methods are insufficient for resolving the complex projection patterns of multiplex projection neurons. Recently MERGE-Seq and Projection-seq, which use a low-cytotoxicity rAAV2-retro virus, have achieved deep analysis of gene profiles with projectome information [21, 22]. Yet, the spatial locations of the neurons and the architecture of the network are still lacking, hampering the comprehensive understanding of the organization principles and wiring logic of neural network.

Prefrontal cortex (PFC) is a critical integration center within the brain network, responsible for a wide range of essential functions including cognition, decision-making, memory, and emotions [23, 24]. Dysfunctions in PFC circuitry and its related functions can lead to various cognitive and neuropsychiatric disorders [25, 26]. Single-cell RNA sequencing and spatial-omics analysis have revealed the sophisticated cellular architecture of the PFC across different species [27–29]. Additionally, the global PFC neural network has been investigated by several technologies such as fMOST and the combination of neural circuit tracing with single-cell sequencing, providing valuable insights into the neural connections between the PFC and other brain regions [22, 30, 31]. However, the multi-modal PFC atlas encompassing neuronal connectivity, transcriptomes, and spatial organization remains fragmented and lacks all-inclusive integration. This hampers the deep analysis of the organizational logic of PFC neural circuits and the biological functions. For instance, how are the diverse neurotransmitter and neuropeptide receptors elegantly configured within distinct circuits to specifically decode the neural input signals? What is the organization and synchronization logic of different projectomic and transcriptomic cell clusters, and the molecular mechanism underlying the wiring of these circuits?

The aim of this study is, therefore, to develop a cost-effective, high-throughput neural circuit tracing method with the robust capacity to simultaneously decipher neural projectome, transcriptome and spatial organization information at single-cell resolution. With the combination of barcoded tracing virus, single-cell sequencing and spatial-omics, we developed a robust Single-cell Projectome-transcriptome In situ Deciphering Sequencing (SPIDER-Seq) method. Leveraging SPIDER-Seq, we delineated an atlas of mouse PFC integrating projectomics, transcriptomics and spatial-omics information (33,766 cells for single-cell sequencing and 124,829 cells for spatial-omics). This publicly available multi-modal dataset (https://huggingface.co/spaces/TigerZheng/SPIDER-web) offers an unprecedented view of the neural circuitry in the PFC and shed deep insights into the neural circuit-specific gene expression pattern, spatial distribution, neural transmission information and neural wiring organizing principles of mouse PFC. Notably, the multi-modal dataset generated by SPIDER-Seq can be trained to predict PFC neuron projections with high accuracy by combining gene profiles and spatial information via machine learning.

## RESULTS

### Deciphering the PFC spatial projectome, transcriptome architecture by SPIDER-Seq

Despite the rapid advance of projectome-transcriptome decoding methods, there are still multiple urgent needs for the improvements in: 1) reducing the cytotoxicity of tracer viruses which could significantly induce gene profile alterations; 2) detecting spatial location and the surrounding local environment information of the neurons in the network at a high resolution; 3) integrating multi information modalities in the same sample; 4) simultaneous high multiplex targets tracing. To help address these requirements, we developed Single-cell Projectome-transcriptome In situ Deciphering Sequencing (SPIDER-Seq), a cost-effective method to achieve high-throughput, single-cell resolution tracing of projection neurons along with transcriptional and spatial profile in mouse PFC (**Fig. 1A**). First, we generated the retrograde rAAV2-retro tracing virus library containing diverse DNA barcodes (**Fig. S1A**), of which the barcode sequences are listed in **Table S1**. To validate the specificity of the rAAV2-retro-barcode virus, we injected the virus into the ventral striatum (ACB) and examined the cellular distribution patterns using fluorescence *in situ* hybridization. As shown in **Fig. S1B**, barcode signals were predominately observed in excitatory neurons (barcode merged with *Slc17a7* positive cell), this is consistent with the view that the long-range projection neurons in the cortex are predominantly excitatory neurons [16]. Moreover, we also validated that one cell can be infected by more than 10 different rAAV2-retro viruses in vitro (data not shown), which was also confirmed by the following in vivo experiments (**Fig. S1C**).

**Fig. 1:**
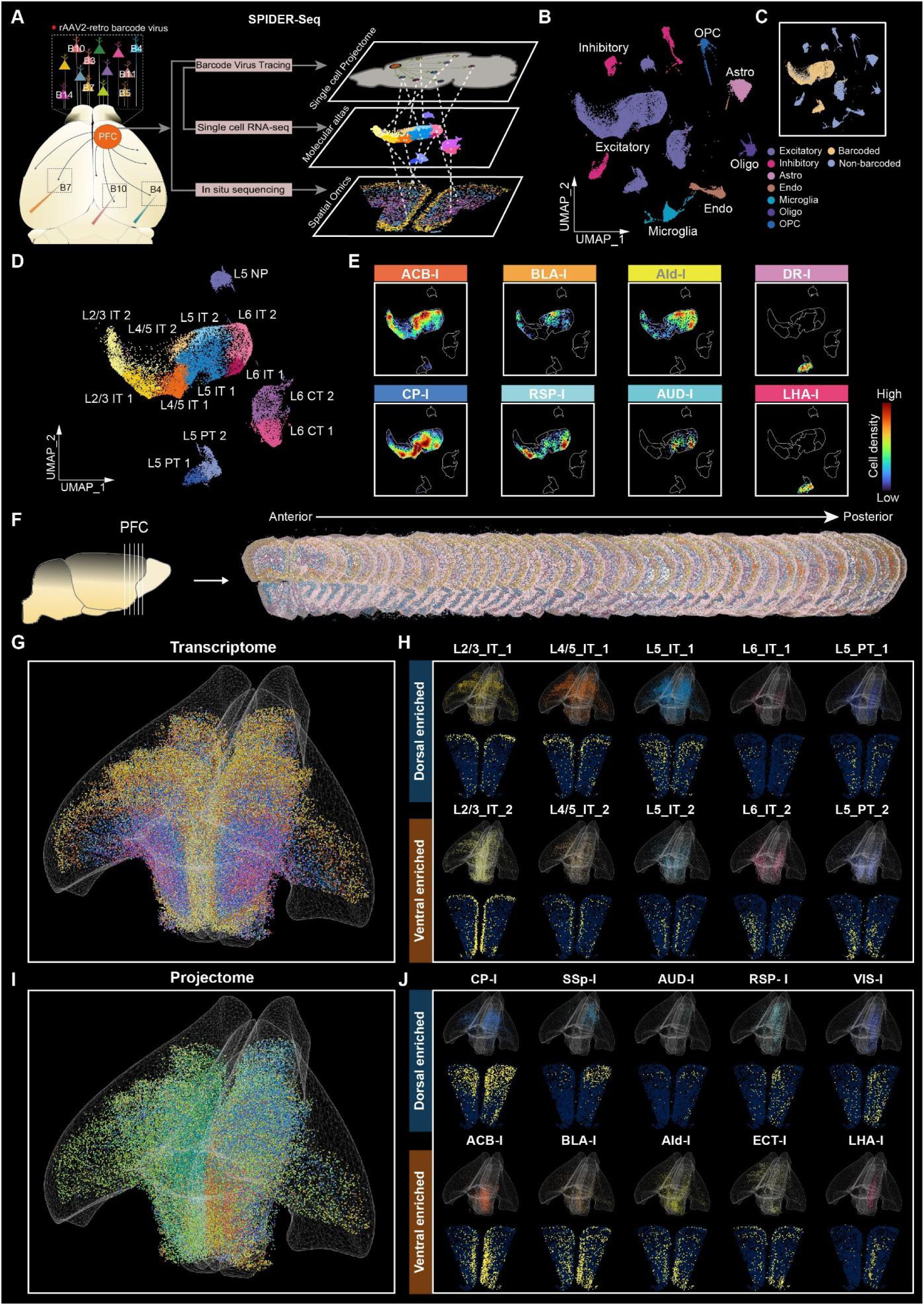
Delineating multi-modal PFC atlas embedding single-cell projectomics, spatial-omics and transcriptomics by SPIDER-Seq. (**A**) Schematic diagram of the SPIDER-Seq workflow. The rAAV2-retro barcode viruses were injected into different nuclei of the mouse brain to achieve high-throughput single-cell resolution retrograde tracing of the PFC projection circuits. 30 days post-injection, single-cell RNA sequencing and *in situ* sequencing were performed on the PFC tissue to decipher the projectome with transcriptomic and spatial information. (**B**) UMAP of all cells in scRNA-seq, colored by main cell clusters. (**C**) Distribution of barcoded cells in UMAP plot. (**D**) UMAP plot shows PFC excitatory neurons clustered into 13 unique subtypes. (**E**) Density scatter visualization on UMAP of neurons projecting to different downstream targets. Points are colored by cell density. (**F**) *In situ* sequencing of barcodes and marker genes of excitatory neuron subtypes in 36 consecutive coronal slices of PFC (Bregma: 2.8mm-0.5mm). (**G**) 3D visualization of the spatial distribution of PFC excitatory neuron subtypes, which displayed separately in (H) and Fig. S1K. (**H**) Spatial distribution of different excitatory neuron subtypes in 3D (top) and 2D (bottom) within an example slice (Bregma: 2.1mm). (**I**) 3D visualization of the spatial distribution of neurons projecting to 15 downstream targets, which displayed separately in (J) and Fig. S1K. (**J**) Spatial distribution of neurons projecting to different targets in 3D (top) and 2D (bottom) within an example slice (Bregma: 2.1mm).

Next, the unique barcoded viruses were injected into 24 main downstream targets of mouse PFC, covering the majority of the whole projection outputs of PFC according to Allen Mouse Brain Connectivity Atlas. After extensive optimization, we achieved precise injections in up to 16 nuclei within the same brain, enabling us to redundantly cover all 24 target nuclei with multiplexed injections of in just three mice (**Table S2**), in which the reproducibility was validated through subsequent SPIDER-Seq analysis. The validation of fluorescence of injection sites was demonstrated in **Fig. S2**. We waited 30 days post-injection to allow the complete expression of the barcode, and then performed single-cell sequencing on the retrogradely labeled PFC tissue. To further delineate the spatial architecture of the PFC transctiptome and projectome, we performed in situ sequencing for 47 sequences including 32 excitatory neuron subtype marker genes and 15 circuit tracing barcodes across 36 continuous slices spanning the entire PFC (**Fig. 1F**). As the marker genes are quite unique or predominantly expressed in different excitatory neurons subtypes, the combination of these marker genes and the barcode information could be enough to differentiate the neurons’ subtypes and align these omics data (**Fig. S5B**). The integrated analysis of multi-modal data from SPIDER-seq provided an atlas with the spatial organization, molecular landscape, anatomy and the projectome information of the projection neurons (33,766 cells for single-cell sequencing and 124,829 cells for spatial-omics) in the PFC at single-cell resolution.

Through single-cell RNA sequencing, 33,766 cells with high quality were clustered into 7 main types (**Fig. 1B** and **S1D-F**). Among these, 18,615 neurons (4,964 genes per neuron) were further classified into 11 types, consistent with previous study [27] (**Fig. S1G**). Next, we conducted elbow analysis to carefully filter out background barcode UMIs (**Fig. S3A** and **D**), and obtained 9,038 barcode-labeled cells projecting to 24 downstream targets (**Fig. 1C** and **S3B-C**). The barcode-labeled cells were predominately excitatory neurons, and its proportion in non-neurons was below 0.5% (**Fig. S1H**), supporting the integrity of our data and the analysis pipeline. Of note, we observed that 67.2% of the neurons target more than one nucleus (**Fig. S1C**), with neuron being co-infected by up to 13 barcoded viruses. This indicated that our approach can be effectively applied for multiplex downstream target nuclei tracing.

Next, the excitatory neurons were further clustered into 13 transcriptomic subtypes. Except for NP (near-projecting) subtypes, neurons in each layer were categorized into two distinct subtypes (**Fig. 1D** and **S1I**), suggesting the presence of transcriptomic differentiation within each layer of PFC neurons. According to the viral barcode information, we mapped the PFC projection neurons targeting to the 24 downstream nuclei onto this detailed transcriptomic atlas, respectively. Our data revealed that the neurons targeting the same downstream nucleus are distributed across multiple transcriptomic subtypes. And the neurons targeting to different downstream nuclei exhibited distinct distributions in the transcriptomic atlas (**Fig. 1E** and **S1J**).

Next, we performed *in situ* sequencing for 47 genes by multiplex detection (**Fig. S4**) based on the modified MiP-seq protocol [32]. Overall, we obtained 124,829 PFC neurons with spatial information, of which 76,512 were barcode-labeled cells. By Tangram analysis [33], we integrated the single-cell RNA sequencing data with the in situ sequencing data, thereby delineating the spatial transcriptomic architecture of the PFC (**Fig. 1G** and **S5A**). The spatial transcriptome data revealed a spatial gradient distribution pattern of the excitatory transcriptomic cell clusters (**Fig. 1H** and **S5B-C**) and the marker genes within each layer of the PFC (**Fig. S5D**). For example, both IT and PT neuron clusters exhibit a dorsal to ventral subtype separation trend in each layer (**Fig. 1H**). Based on this spatial transcriptomic information, we categorized PFC IT and PT neurons into dorsal-enriched and ventral-enriched cell clusters, respectively (**Fig. 1H**). Meanwhile, we constructed a three-dimensional spatial projectome map of the projection neurons targeting 15 downstream targets in the same mouse with single-cell resolution (**Fig. 1I-J** and **S1K**). Notably, the spatial projectome also reveal a similar dorsal-ventral distribution trend, suggesting a potential interlink between the transcriptome and projectome in the PFC.

To validate the accuracy of in situ sequencing, we first compared the marker gene distribution patterns of our data with Allen ISH atlas, which showed consistent results (**Fig. S5C**). Then we compared the distribution of viral barcode signal obtained by *in situ* sequencing with the florescencent rAAV2-retro tracing results in indpendent mice, and the repeat measurements in another mouse. The consistent distribution patterns in **Fig. S6** demonstrated the accuracy and reproducibility of our barcodes detection.

Meanwhile, as the barcode signal retrogradely traced from LHA, one of the PT neuron projecting targets, should located in layer L5, we validate its colocalization with layer L5 pyramidal neuron specific marker gene *Pou3f1* by FISH assay. As shown in **Fig. S7**, the barcode signals that retrogradely traced from LHA is indeed mostly merged with *Pou3f1* in layer L5, supporting the specificity integrity of our experiments.

Next, we conducted more systematic quality control, reproducibility and integrity analysis of SPIDER-Seq. First, we compared the differences between barcoded and non-barcoded neurons to evaluate the impact of viral infection on gene expression of the infected neurons. The Pearson correlation between these two groups is 0.98 (**Fig. S1L**), indicating that the effect of rAAV2-retro virus infection on gene expression is negligible. Importantly, we also compared the projection patterns and intensity between different replicates resolved by SPIDER-Seq, yielding an overall Pearson correlation between different samples of 0.81 (**Fig. S1M**), which supported the reproducibility of our viral infections and data interpretation. Moreover, we compared the projection profiles resolved by single-cell sequencing and in situ sequencing, revealing a Pearson correlation of 0.80 (**Fig. S1N**), further demonstrating the reproducibility between these two independent approaches. Additionally, we validated our PFC projection profiles resolved by SPIDER-Seq with the fMOST data from Gao’s study (**Tables S6**) [31], achieving a Pearson correlation of 0.81 (**Fig. S1O**). Together, these data demonstrated the reproducibility and integrity of our SPIDER-Seq data. This robust method may greatly facilitate to delineating the brain multi-modal network atlas and understanding neural circuit organization logic.

### Spatial and transcriptomic configuration of PFC projectome by SPIDER-Seq

The SPIDER-seq multi-modal PFC atlas embedding large scale neural circuit, transcriptomes, and spatial architecture information at single-cell resolution provides a unique opportunity to understand the underlying cellular and circuital organization logic. To interpret the relationship among projectome, transcriptome, anatomy and the spatial organization principles, we first analyzed the architecture of the projection neurons. We observed that neurons targeting different downstream nuclei exhibit unique spatial distributions. For example, neurons targeting SSp-I, AUD-I, RSP-I and VIS-I are predominantly located in the posterior dorsal part of PFC, whereas neurons targeting ECT-I, AId-I, BLA-I and ACB-I are found mainly in the anterior ventral part (**Fig. 2A** and **B**). As illustrated in **Fig. 2C-D** and **S8A**, the projection neurons targeting to ACB-I and SSp-I gradiently occupied different territories in the 3D space of PFC and consist of different transcriptomic cell subtypes, respectively. These data may illuminate how the projection neurons synchronously configurate their soma (for transcriptome) and axon (for projectome) in the PFC. The 3D atlas of the projectome targeting each nucleus together with the transcriptome of each excitatory subtype were shown in **Supplementary Video 1**. These data suggested that the spatial transcriptome and projectome were geographically configured in PFC with distinct organization principle.

**Fig. 2:**
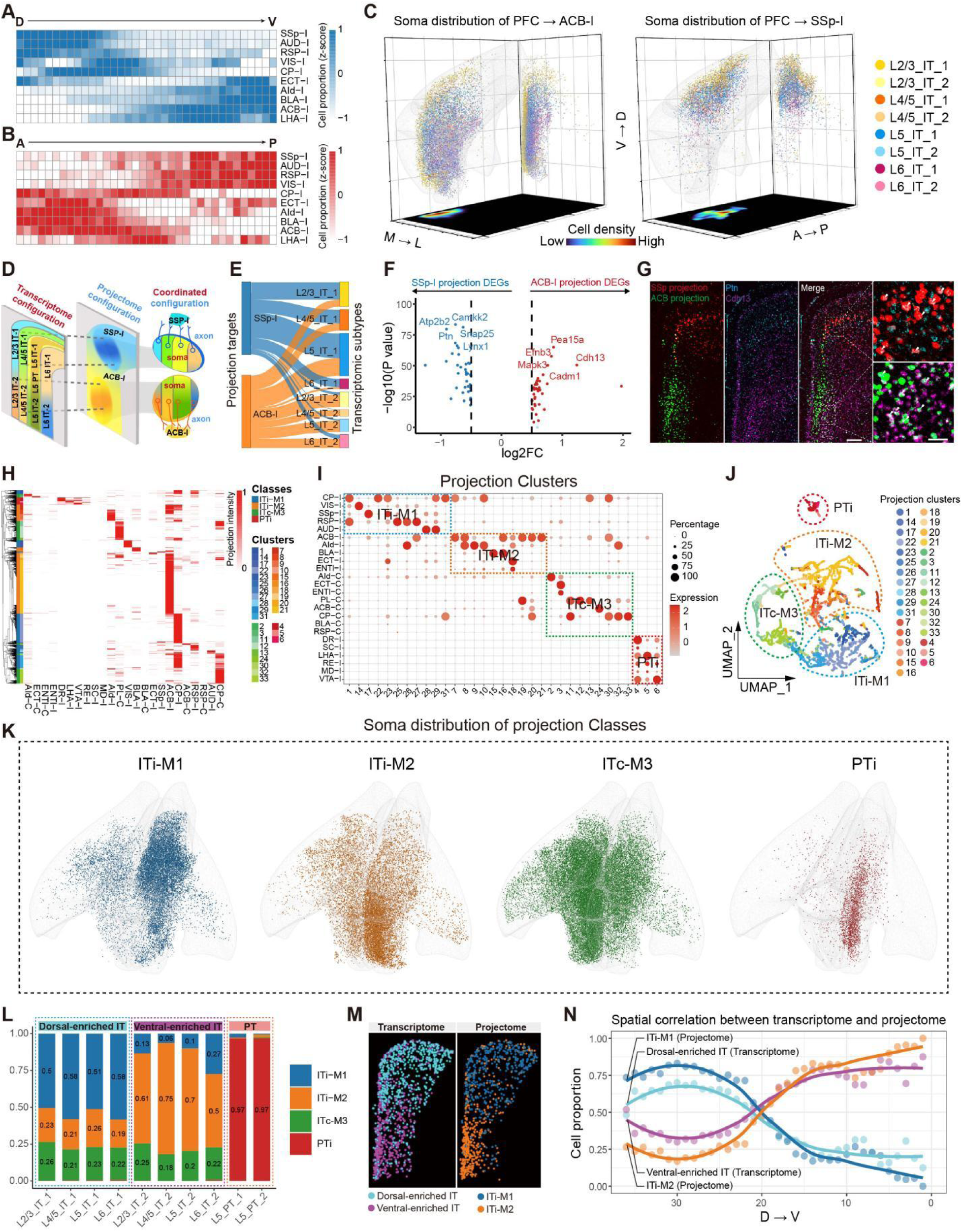
Deciphering the spatial and transcriptomic configuration of PFC projectome by SPIDER-Seq. (**A**) Heatmap showing the proportion of PFC neurons projecting to different downstream nuclei from dorsal to ventral (D to V) axis. The cell proportions were normalized by row to calculate the z-score. (**B**) Heatmap showing the proportion of PFC neurons projecting to different downstream nuclei from anterior to posterior (A to P) axis. The cell proportions were normalized by row to calculate the z-score. (**C**) 3D visualization of the spatial distribution of PFC neurons projecting to ipsilateral ACB-I and SSp-I, colored by transcriptomic subtypes. Right grid shows all neurons superimposed on the coronal plane. Bottom grid shows the density of all neurons on the transverse plane. (**D**) Schematic diagram showing the layered distribution of PFC transcriptome and the spatial gradient organization across layers of projectome. (**E**) Sankey diagram showing the transcriptomic subtypes composition of PFC neurons projecting to ACB-I and SSp-I, respectively. (**F**) Differentially expressed genes (DEGs) between ACB-I and SSp-I projection neurons. (**G**) *In situ* hybridization assay showing that the ACB-I enriched gene *Cdh13* accumulated in the ventral PFC, whereas the SSp-I enriched gene *Ptn* accumulated in the dorsal PFC (Bregma: 2.1mm). Blue, *Ptn*, purple, *Cdh13*, red, SSp-I, green, ACB-I. Scale bar: 500 µm. Magnified view of the white boxed area (right). Scale bar: 50 µm. (**H**) Heatmap of the projection intensity of 9,038 PFC projection neurons to 24 targets. Each row represents the projection intensity of a single neuron. Neurons are divided into four projection classes and 33 projection clusters by hierarchical clustering. (**I**) Dot plot shows the projection of 33 projection clusters to 24 targets. The color and size represent projection intensity zscore and cell percentage, respectively. The dashed box highlights four projection classes. (**J**) UMAP analysis of the PFC projection matrix showing the distribution of 33 projection clusters. These projection clusters can be divided into four projection classes: PTi, ITi-M1, ITi-M2, and ITc-M3. (**K**) 3D visualization of the spatial distribution of four projection classes in PFC. (**L**) The transcriptomic subtypes proportion of the four projection classes, colored by projection classes. (**M**) Spatial distribution of dorsal-enriched IT and ventral-enriched IT transcriptomic subtypes (left). Spatial distribution of ITi-M1 and ITi-M2 projection classes (right) (Bregma: 2.1mm). (**N**) Spatial correlation between transcriptome and projectome of IT neurons. The color of lines corresponds to (M).

With the integrated transcriptome information, SPIDER-Seq can also reveal the transcriptomic differences between various types of projection neurons. For example, the neurons targeting SSp-I and ACB-I not only exhibit different spatial distribution, but also show distinct transcriptomic subtype compositions. Neurons from L2/3IT1, L4/5IT1, and L6IT1 predominantly target SSp-I, while those from L2/3IT2, L4/5IT2, L5IT2 and L6IT2 prefer to target ACB-I (**Fig. 2E**). Additionally, the SSp-I projecting neurons express significantly high levels of *Snap25*, *Camkk2*, and *Ptn*, while the ACB-I projecting neurons highly express *Pea15a*, *Efnb3*, and *Cdh13*, which was validated by FISH assays (**Fig. 2F** and **G**).

We also observed specific gradient spatial distribution patterns of neurons projecting to different regions of striatum. Neurons projecting to the ipsilateral ventral striatum (ACB-I) primarily located in the ventral region (**Fig. 2C** and **S8C**), while those projecting to ipsilateral dorsal striatum (CP-I) tend to concentrate in the dorsal region (**Fig. S8B** and **C**), which is consistent with a previous study [34]. Meanwhile, we found transcriptomic differences between the neurons targeting the ventral and the dorsal striatum as shown in **Fig. S8D**. These data highlighted the high throughput capability of SPIDER-Seq to systematically analyze neuronal projection patterns, gene expression as well as spatial organization at single-cell level.

To further analyze the detailed projection patterns of PFC neurons, we performed hierarchical clustering on the single-cell projection profile of each neuron, resulting in the identification of 33 distinct projection clusters (**Fig. 2H-J**). These projection cell clusters can be grouped into four projection classes: PTi, ITi-M1, ITi-M2 and ITc-M3, each displaying distinct three-dimensional spatial distribution patterns within the PFC (**Fig. 2K** and **S8E-F**). PTi were clustered into a separated class, sending projection to subcortical regions such as LHA-I, VTA-I, DR-I et al. IT neurons were further characterized based on their projection patterns as follows: (1) ITi-M1 class mainly project to the ipsilateral dorsal striatum (CP-I), medial and dorsal cortical regions including RSP-I, VIS-I, AUD-I, and SSp-I, of which the somas located in the dorsal part of the PFC. Anatomically, ITi-M1 mostly restricted to ACC; (2) ITi-M2 class primarily project to the ipsilateral ventral striatum (ACB-I), BLA-I, and lateral cortical regions including AId-I, ECT-I, and ENTI-I, of which the somas located in the ventral part of the PFC, around PL and IL anatomic regions; (3) ITc-M3 neurons primarily project to the contralateral brain regions including AId-C, ECT-C, ENTI-C, PL-C, ACB-C, and CP-C (**Fig. S8G**).

Next, we performed spatial projectome and transcriptome integrated analysis on these projection classes and clusters, and identified distinct differentially expressed genes in each clusters (**Fig. S8H-J**). Consistent with previous data [17], neurons in the PTi class are pyramidal tract neurons originating from layer 5, largely composed of L5-PT-1/2 transcriptomic clusters (**Fig. 2L**). In contrast, the IT classes comprises neurons from various transcriptomic clusters across different layers (**Fig. 2L**). While most projection clusters contain neurons from multiple layers, several projection clusters exhibit strong preferences for their transcriptomic cell type composition. For example, 75.0% neurons of projection cluster 3, which simultaneously targets ECT-C, AId-C, and LENTI-C, belong to L2/3-IT transcriptomic clusters. Similarly, 69.2% neurons of the projection clusters 26 targeting RSP-I and AId-I consist of L6-IT transcriptomic clusters (**Fig. S8K** and **L**). The identification of the projection specific transcriptomic signature genes would help PFC researchers develop new genetic tools to target and manipulate those cells to better understand the functions of specific circuit.

We also observed transcriptomic differences between ITi-M1 and ITi-M2 classes. ITi-M1 projecting neurons are primarily composed of the dorsal-enriched transcriptomic clusters, while ITi-M2 projecting neurons are mainly derived from the ventral-enriched transcriptomic clusters (**Fig. 2L**). Notably, the spatial distribution of ITi-M1 and ITi-M2 projection classes is highly correlated with spatial gradients of the dorsal- and ventral-enriched transcriptomic cell clusters (**Fig. 2M** and **N**), suggesting a synchronized configuration of different neuron projection clusters and transcriptomic cell types in the PFC.

### Spatial, anatomic and trascriptomic configuration of PFC IT projection neurons

Given the high diversity in the projections of IT neurons, we further analyzed the spatial and cellular configuration principles of PFC IT projection neurons. To quantify the diversity of IT projection patterns, we binarized the projection to downstream targets, and presented the landscape of the transcriptomic cell type composition and spatial information of the top 50 IT projection motifs (**Fig. 3A**). We exemplified five projection motifs targeting AId-I to demonstrate their detailed spatial, transcriptomic, and anatomic configuration. Different projection motifs targeting AId-I exhibit a gradient spatial transition from dorsal to ventral regions correlated to the axon branching locations in striatum (ACB-I or CP-I) (**Fig. 3B**). Neurons projecting to both AId-I and ACB-I are mainly located in the ventral part of the PFC, while those targeting both AId-I and CP-I predominantly distribute in the dorsal part of the PFC, and the triple-targeting neurons (AId-I, CP-I and ACB-I) located in an intermediate zone, with distinct transcriptomic signatures respectively (**Fig. 3B** and **S9A**). Similar patterns also appear in the BLA-I targeting projection motifs, as shown in **Fig. S9B** and **C**.

**Fig. 3:**
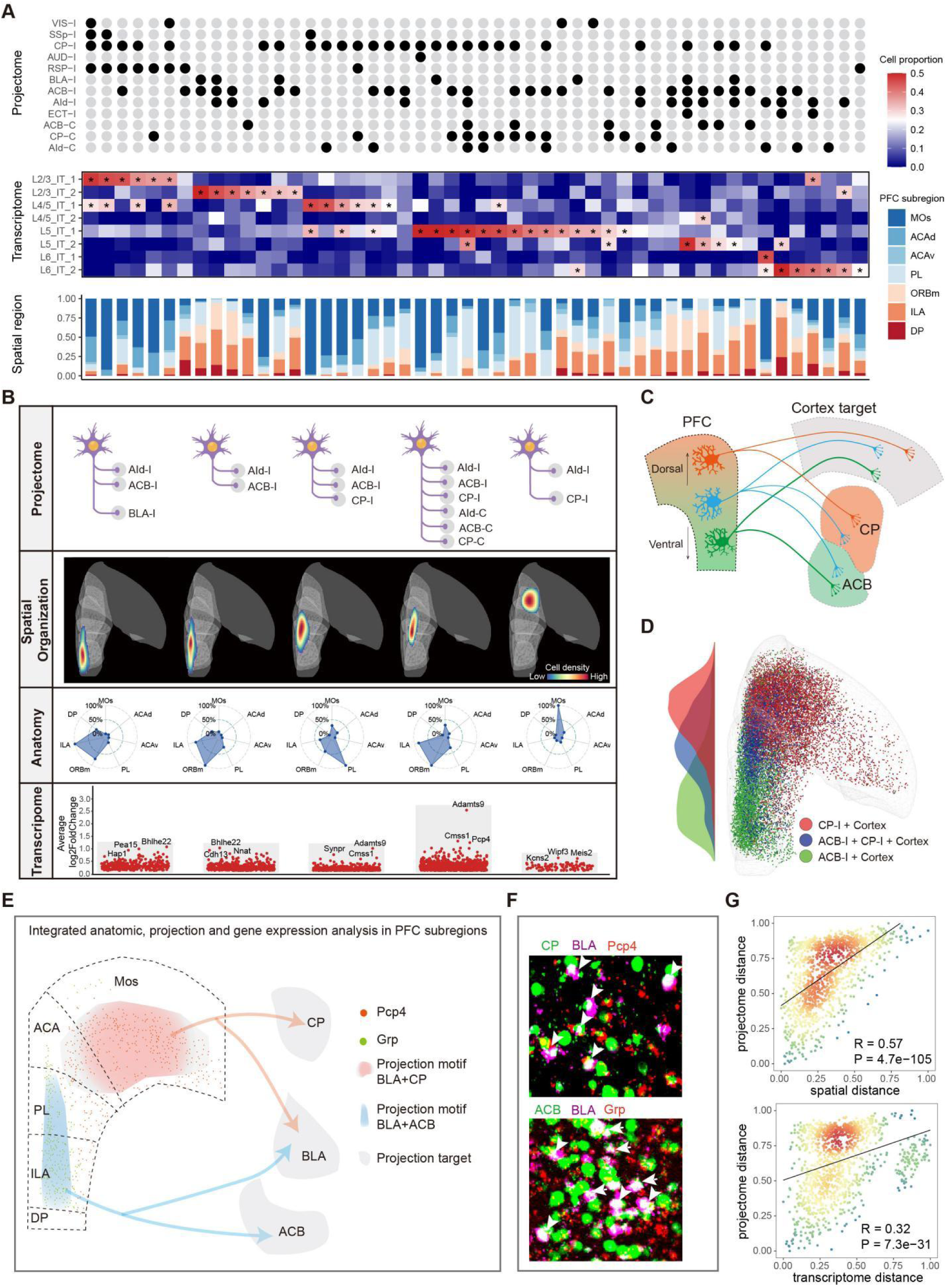
Integrative analysis of spatial and transcriptomic organization pattern of PFC IT projection neurons. (**A**) Integrative analysis of PFC IT projection motifs, transcriptome subtypes, and spatial distribution. The top 50 projection motifs based on binarized projection matrix (top), transcriptomic subtypes (middle), and spatial distribution in different PFC anatomical subregions (bottom) of individual projection motifs. Asterisks: cell proportion larger than 0.25. (**B**) Transcriptional, anatomic and spatial information of different PFC IT projection motifs targeting AId-I. Top, projection motifs. Upper middle, spatial distribution. Lower middle, anatomy composition. Bottom, differentially-expressed genes (DEGs) of 5 AId-I projection motifs. (**C**) A schematic diagram showing the spatial distribution of the neurons co-projecting to cortex (or BLA) and striatum (CP or ACB or both). (**D**) Spatial distribution of neurons with CP-I+Cortex (or BLA) (red), ACB-I+Cortex (or BLA) (blue), and ACB-I+CP-I+Cortex (or BLA) (green) projection motifs. (**E**) Integrated anatomic, projection and gene expression analysis in PFC subregions. The distribution of two projection motifs targeting BLA is consistent with DEGs. (**F**) Barcode neurons of two projection motifs (BLA+CP, BLA+ACB) merged with DEGs. (**G**) Correlation between projectome and spatial location (top) or transcriptome (bottom). Each point represents the spatial or transcriptome euclidean distance (x-axis) and projection euclidean distance (y-axis) between a pair of projection motifs in (A), colored by the scatter density.

To further test whether this pattern represents a common organization principle, we systematically analyzed the spatial architecture of all the cortex and striatum co-targeting IT neurons in the PFC. In our dataset, 84% of IT neurons that project to the cortex/BLA also extend their axons to striatum (CP or ACB) (**Fig. S9D**). These neurons exhibit distinct spatial distributions and spatial transcriptomic signatures depending on their targeting in striatum. The projection neurons co-targeting dorsal striatum (CP) are mainly distributed in the dorsal part of the PFC, while those co-targeting ventral striatum (ACB) are primarily enriched in the ventral part. Projection neurons that simultaneously targeting both ACB and CP are predominantly located in the intermediate transitional zones (**Fig. 3C** and **D**). Correspondingly, these projection neurons showed distinct gene profiles, of which some transcriptomic signatures also display a dorsal-enriched to ventral-enriched gradient, as confirmed by FISH assay (**Fig. S9E-G**).

We also exemplified an integrated anatomy, projection and transcriptomic analysis and identified the CP and BLA projection neurons are mainly located in Mos regions in PFC highly expressing *Pcp4*, while the ACB and BLA projection neurons are mainly located in PL and ILA regions in PFC highly expressing *Grp*, which was further confirmed by FISH assay (**Fig. 3E** and **F**). Importantly, our analysis revealed significant correlations between neuron projections and their gene profiles, and spatial location, respectively (**Fig. 3G**). Together, these data suggest that although the projectomic and transcriptomic cell clusters display distinct spatial organization principle, they are synchronously configured in the PFC. Our analysis could provide unprecedented integrated anatomic and transcriptomic formation at projection motif levels, which may greatly contribute to understanding the function of each anatomic regions in PFC.

### Co-projection principle of PFC IT neurons

Our data indicated that the majority of barcoded neurons target two or more nuclei simultaneously (**Fig. S1C**). To examine whether these downstream targets are randomly associated together or sophisticatedly organized with certain kinds of logic, we aligned the projection patterns of multiple targeting neurons resolved by SPIDER-Seq with the expectations of random association (**Fig. 4A** and **S10A**). The results suggested that these multiple-projections are not randomly configurated. For example, the multiple-projection neurons targeting both ACB-I and BLA-I are overrepresented, whereas the multiple-projection neurons targeting CP-I and AId-I are underrepresented (**Fig. 4B** and **S10B**).

**Fig. 4:**
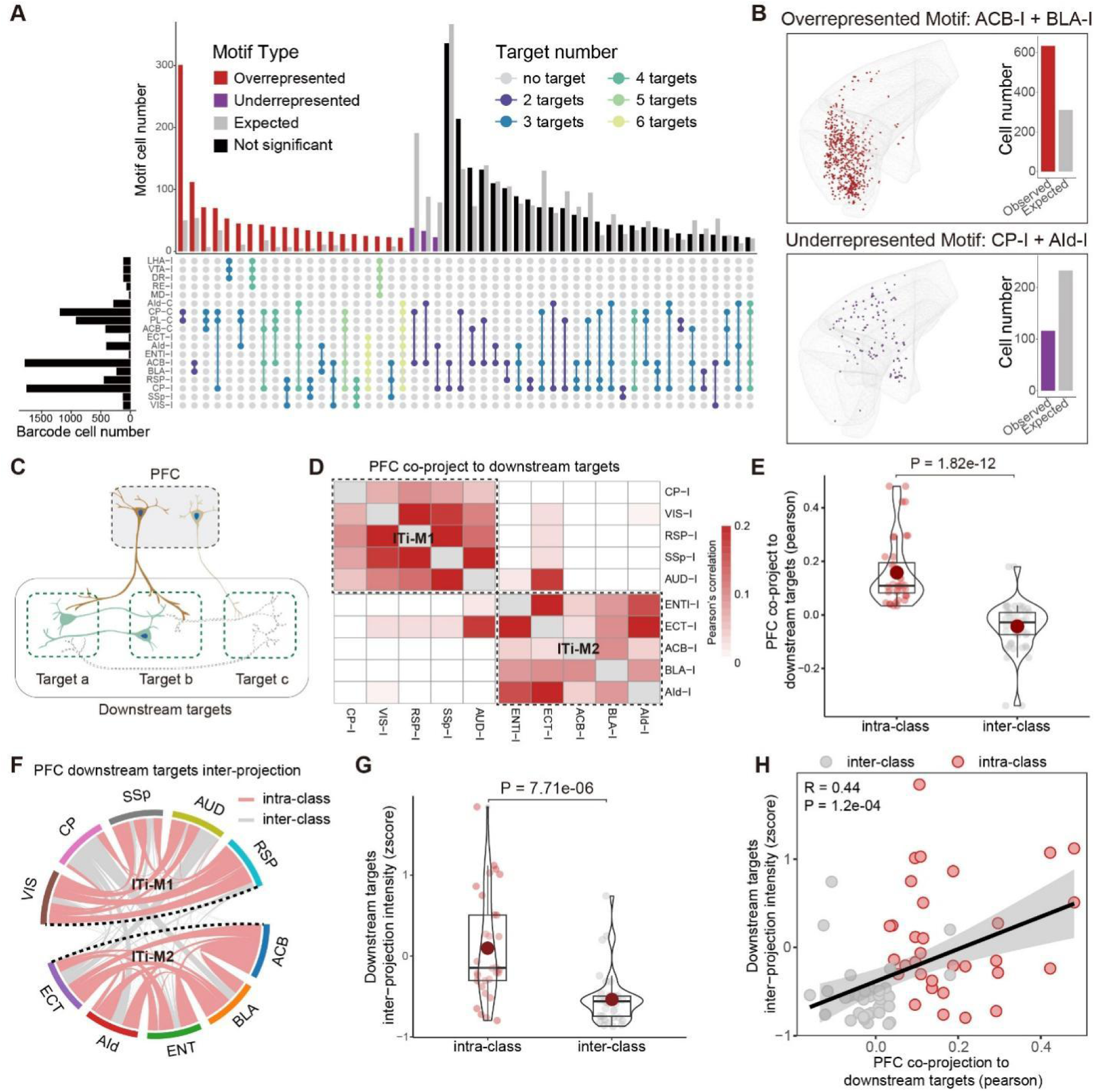
Wiring logic analysis of PFC IT co-projection neuron. (**A**) Upset plot showing the observed and expected cell numbers for each projection motif. We compared the observed cell numbers of projection motifs with the expected null model to calculate significance. Red, purple, and black bars represent overrepresented, underrepresented, and not significant, respectively. Gray bars represent expected. Different projection motifs are colored according to the target number. (**B**) Example of spatial distribution of overrepresented (ACB-I+BLA-I) and underrepresented (CP-I + AId-I) motifs. The barplot showing the observed and expected cell numbers for motifs in spatial dataset. (**C**) A schematic diagram showing that the downstream targets co-projected by PFC IT neuron share more circuit connections. (**D**) Heatmap showing the co-projection probability of downstream targets targeted by PFC IT neuron. Each tile represents the pearson correlation of two targets co-projected by PFC projection neurons. (**E**) Box and violin plot showing that the co-projection probability within projection classes is significantly higher than that across classes. (**F**) Circuit connection intensity between downstream targets targeted by PFC IT neurons, connectivity intensity refers to the density of AAV-labeled nerve from Allen Mouse Brain Connectivity Atlas. The red arcs represent intra-class projection, and the grey arcs represent inter-class projection. (**G**) Box and violin plot showing that circuit connection intensity between downstream targets within the same projection classes is higher than that across classes. (**H**) Correlation between co-projection probability and circuit connection intensity between downstream targets targeted by PFC IT neurons.

Furthermore, we investigated the wiring principle of the multi-projection neurons in PFC. First, we analyzed co-projection probability to the downstream targets from different PFC projection neurons. The results showed that downstream targets belonging to the same projection class (ITi-M1 or ITi-M2) have significantly higher probability to be co-projected compared to the targets from different classes (**Fig. 4D** and **E**). Considering the high co-projection probability may accompany the engagement of more simultaneous information processing and the subsequent information exchange, the co-projected downstream targets should presumably have more reciprocal circuit connections (**Fig. 4C**). Thus, we collected the projection data for the corresponding nuclei from the Allen Mouse Brain Connectivity Atlas and calculated the connectivity intensity between the downstream targets (**Tables S7**). The results verified that the downstream targets from the same projection class indeed have significantly more intensive circuit connections (**Fig. 4F-G** and **S10C-D**). Additionally, there is a positive correlation between the connectivity intensity of downstream nuclei and the probability of co-projecting by PFC neurons (**Fig. 4H**). These findings revealed one wiring principle in PFC and validated that projection of the PFC neurons are indeed sophisticatedly organized rather than randomly wired together.

### Configuration of neural signal decoding and transmission machineries in the content of PFC neural network

PFC neurons decode upstream information inputs through the neurotransmitters and neuropeptides receptors, transmitting signals to downstream networks by releasing neurotransmitters and neuropeptides. Deciphering the dynamic and sophisticated neural signal transmission flow in neural network is the prerequisite for the understanding of neural computation and brain functions [35, 36]. However, the detailed signal decoding and transmission processes remain elusive. Our SPIDER-Seq dataset contains high quality of gene profiles data which reach to 4964 genes per neuron and have important expression information including low-expression genes such as the receptor genes for neurotransmitters and neuropeptides. This empowered us to systematically delineate the expression patterns of the various neural signaling related molecules genes in the content of PFC network, which could bring unprecedented insights into the logic of neural signal encoding in different circuits.

Hence, we depicted the neural signaling molecules heatmaps for different projection clusters based on the expression of neurotransmitter transporters, neuropeptide precursors, and their receptor genes (**Fig. 5A** and **S11A**). The heatmaps revealed significant differences in the expression of neural signaling molecules between PT and IT projection neurons, including glutamate receptors (*Gria4*, *Grm3*), serotonin receptors (*Htr2a*, *Htr5a*, *Htr1b*), dopamine receptors (*Drd1*), various neuropeptide receptors (*Mc4r*, *Mchr1*, *Npr3*, *Cckbr*), as well as neuropeptide precursors (*Pdyn*, *Adcyap1*, *Grp*, *Npy*, *Penk*, *Cck*) and neurotransmitter transporter (*Slc17a6*) genes (**Fig. S11B**). These differences may underly the distinct functions of these two types of PFC neurons. Next, we analyzed the neural signaling molecular expression pattern of two IT projection classes targeting the ipsilateral brain areas. Our data showed that the neurons projecting to the dorsal cortical regions / dorsal striatum and those projecting to the lateral cortical regions / ventral striatum showed distinct gene expression patterns of neural signaling molecules (**Fig. 5B**). In this line, previous studies have demonstrated the functional separation of PFC projections to the dorsal medial and the ventrolateral cortex as well as dorsal and ventral striatum [24, 34].

**Fig. 5:**
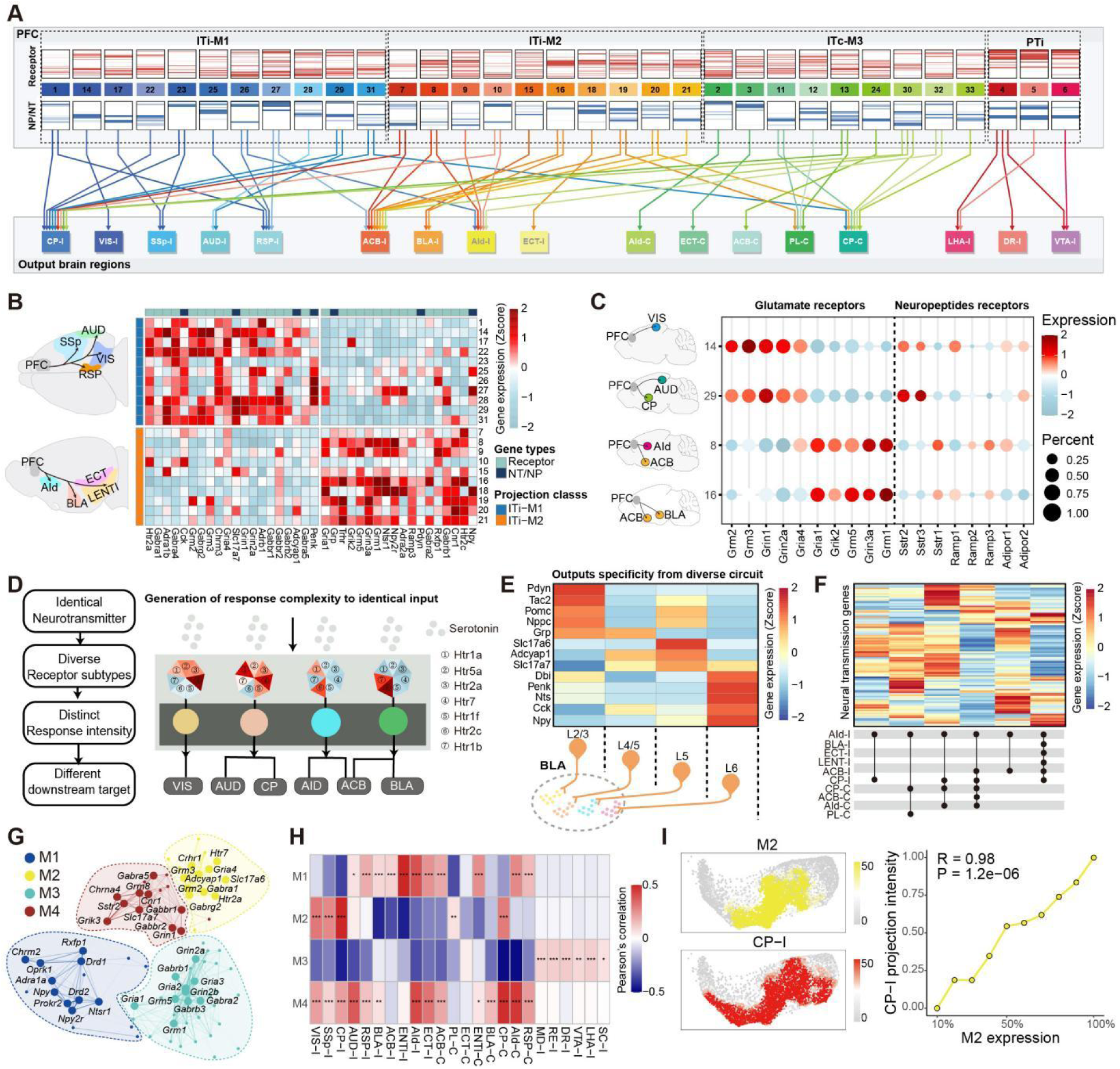
Configuration of neural signal decoding and transmission machineries in the content of PFC neural network. (**A**) Neural signal decoding machineries and transmission flow in PFC projection clusters. The upper panel showing 33 projection clusters grouped by four projection classes. Heatmaps showing the expression of the genes encoding neurotransmitters (NT) (blue), neuropeptides (NP) (blue) and their receptors (red) in each projection cluster. The lower panel showing the downstream targets. Arrows represent the output of each PFC projection cluster to different downstream targets, colored by projection clusters. (**B**) The heatmap displays the differential expression of neural signaling molecules in two projection classes of IT neurons targeting the ipsilateral brain regions (ITi-M1 and ITi-M2). (**C**) Different projection clusters expressing diverse neurotransmitter (glutamate) and neuropeptide (somatostatin, SST; adrenomedullin, ADM; Adiponectin, ADIPOQ) receptor subtypes. (**D**) Schematic diagram showing PFC projection clusters decode serotonin signals by different expression level and combinations receptor subtypes. (**E**) Neurotransmitter transporter and neuropeptide precursor genes expression in different layers neurons targeting the BLA-I. (**F**) Different projection motifs targeting to AId-I have different neural signaling molecules expression patterns. (**G**) Co-expression network of genes related to neural signaling molecules. Each node represents a single gene, and edges represent co-expression links between genes. Genes are divided into four co-expression modules. The top 10 hub genes per module are labeled. (**H**) Heatmap showing the correlation between projection and different gene co-expression modules. Each tile in heatmap represents the pearson correlation coefficient between the projection intensity of a target and the expression of a co-expression module. Statistical significance was determined using two-sided Fisher’s exact test, *P < 0.05, **P < 0.01, ***P < 0.001. (**I**) An example showing that gene co-expression module M2 and projection to CP-I share similar distributions on transcriptome UMAP (left). The correlation between M2 expression and CP-I projection intensity (right).

As the different expression levels and combinations of neurotransmitter receptor subtypes may reflect the differential decoding of the same neurotransmitter input [37, 38], we then examined the expression patterns of neurotransmitter receptor subtypes in different projection clusters. As shown in **Fig. 5C**, glutamate receptor subtypes are distinctly expressed across different projection clusters. The dorsal cortical regions targeting projection clusters, such as cluster 14 (mainly targeting VIS) and 29 (mainly targeting AUD and CP), showing significantly higher expression level of *Grm2*, *Grm3*, *Grin1*, *Grin2a*, and *Gria4*. In contrast, the projection cluster 8 (mainly targeting AId and ACB) and projection cluster 16 (mainly targeting BLA and ACB), highly express *Gria1*, *Grik2*, *Grm5*, *Gria3a*, and *Grm1*. This phenomenon has been also observed in other neuromodulatory receptors, such as neuropeptide receptors (**Fig. 5C**), and serotonin receptors (**Fig. S11C**).

The neighborhood neurons distributed in the same spatial territory in the PFC may likely receive the same local inputs, but could project to different downstream targets. Thus, it would be important to distinctly encode the same inputs into more diverse signals via the combination of differentially expressed signal decoding receptor genes in different projection motifs. By this principle, an identical input has the capacity to encode diverse kinds of downstream signals in the neural network, which may contribute to the generation of enough signal complexity to match the sophisticated information transmission capacity in the brain (**Fig. 5D**).

The release of neurotransmitters and neuropeptides is the primary means by which neurons transmit signals [39]. We observed that the mRNA expression levels of different neurotransmitter transporter and neuropeptide precursor genes vary across different projection clusters (**Fig. 5B**). Notably, projection neurons in distinct layer in PFC targeting the same nucleus show significant differences in the expression of neurotransmitter transporters and neuropeptide related genes (**Fig. 5E**). For example, IT projection neurons targeting BLA from L2/3 highly express *Pdyn* and *Tac2*, neurons from L5 highly express *Adcyap1*, *Slc17a7* and *Slc17a6*, whereas neurons from L6 highly express *Npy* and *Penk*. The molecular configuration of this kind of expression pattern may allow the same down-stream target nucleus to unambiguously differentiate inputs from different upstream nuclei, which might be important to guarantee the specificity of the neurotransmission in the intricate neural network.

Furthermore, we analyzed the expression of neural signaling molecules including neurotransmitter transporters, neuropeptide precursors and the related receptor genes in six projection motifs targeting AId-I and observed significant differences in their expression patterns in each projection motif (**Fig. 5F**). Similar patterns also appeared in the RSP-I targeting projection motifs, as shown in **Fig. S11D**. These data suggest that diverse PFC projection circuits are equipped with differential neural signal decoding and transmission machineries, thereby facilitating their functional diversity. We then investigated whether there are co-expression patterns of the neural transmission molecules in different PFC neural projection clusters. By co-expression analysis, we identified four co-expression modules of the neural signaling related molecules (**Fig. 5G** and **S11E-G**). Importantly, our data revealed a high correlation between the projection patterns and the co-expression of neural signaling molecules (**Fig. 5H**). For example, the projection intensity of neurons targeting CP-I is positively correlated with the expression intensity of co-expression module M2 (**Fig. 5I**). These results suggested that the expression pattern of diverse neural signaling molecules in different PFC circuits is not random organized; rather, it forms sophisticated and specific co-expression modules in different circuits, underlying the synergistic action among different neural signaling molecules.

### Correlation of neural circuit wiring molecules with projection patterns

Although the expression of wiring-related genes during development is crucial for neural circuit formation, the expression of specific genes (such as cadherins and axon guidance molecules) in adulthood is critically important for circuit maintenance. To explore the relationship between projection patterns and these molecules, we further analyzed the high-quality transcriptome profiles of PFC projection neurons in content of neural circuit. Our data revealed that different projection classes exhibited distinct expression patterns of neural circuit wiring molecules (**Fig. 6A** and **S12A**). For example, *Igfbp4*, *Rgma*, and *Fam19a1* are highly expressed in PT neurons (**Fig. 6B** and **C**). In addition to the significant differences between PT and IT projection neurons, projection neurons within different IT classes, such as ITi-M1 and ITi-M2, also showed differential gene expression (**Fig. 6A**). For instance, *Cadm2*, *Sema7a*, and *Pcdh7* are highly expressed in the ITi-M1 class, whereas *Efnb3*, *Cdh13*, and *Nov* are highly expressed in the ITi-M2 class (**Fig. 6B** and **C**). Importantly, the expression of these wiring molecules is in proportion to the intensity of the projection classes (**Fig. 6D**).

**Figure 6.**
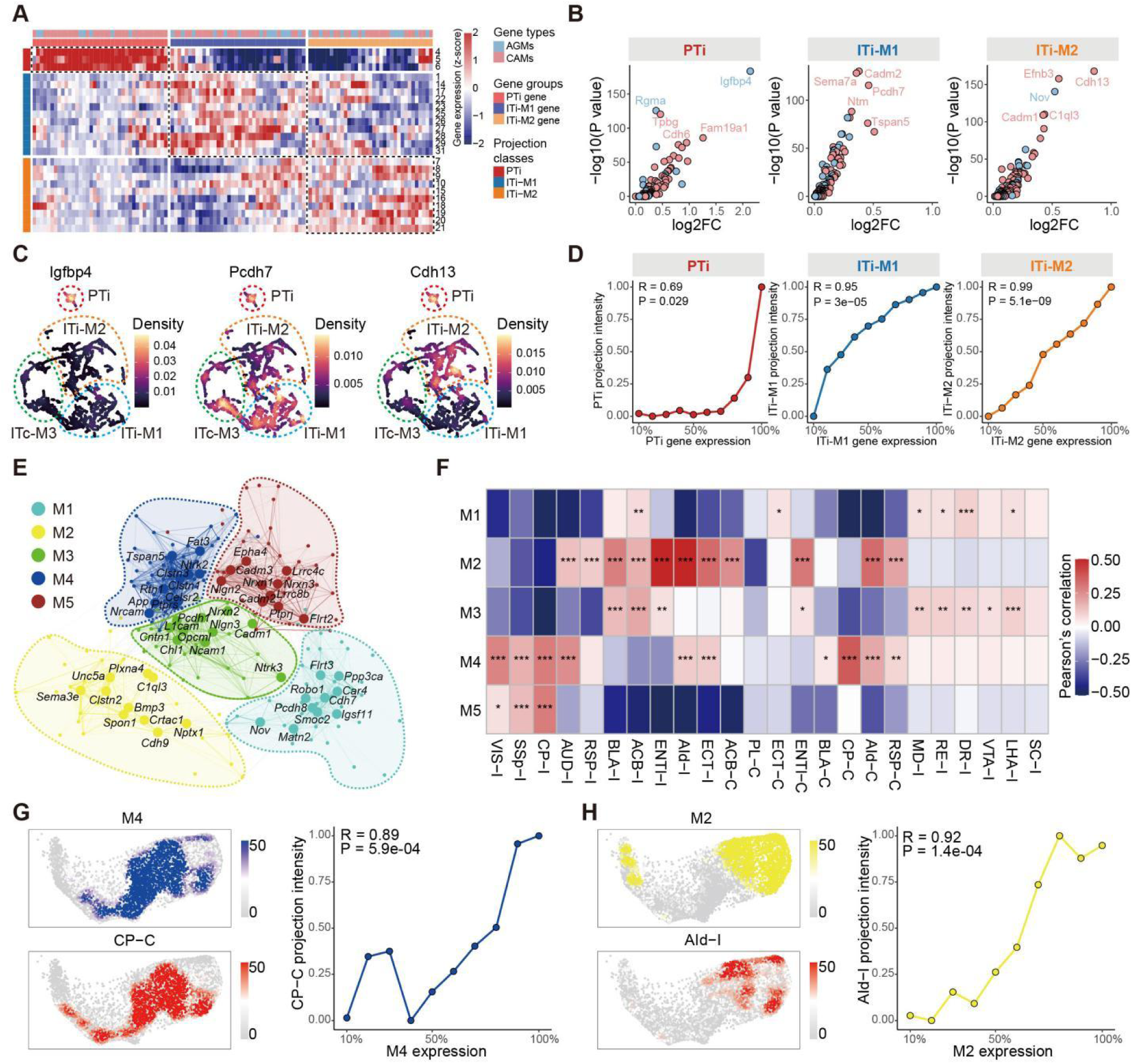
Correlation between expression of neural circuit wiring molecules and PFC projection patterns. **(A)** Expression of molecules related to neuronal circuit wiring molecules (AGMs: axon guidance molecules, CAMs: cadherin molecules) in different projection clusters. The dotted boxes outline the gene groups associated with projection classes. **(B)** Volcano plot showing genes related to neural circuit wiring enriched in PTi, ITi-M1, and ITi-M2 projection classes, respectively. **(C)** Projection UMAP shows the expression distribution of three genes related to neuronal circuit wiring (*Igfbp4* enriched in PTi, *Pcdh7* enriched in ITi-M1 and *Cdh13* enriched in ITi-M2). **(D)** Correlations between gene expression of three gene groups and PTi, ITi-M1, and ITi-M2 projection classes, respectively. **(E)** Co-expression network of genes related to circuit wiring. Each node represents a single gene, and edges represent co-expression links between genes. Genes are divided into five co-expression modules. The top 10 hub genes per module are labeled. **(F)** Heatmap showing the correlation between projection and different gene co-expression modules. Each tile in heatmap represents the pearson correlation coefficient between the projection intensity of a target and the module eigengenes (MEs) of a co-expression module. Statistical significance was determined using two-sided Fisher’s exact test, *P < 0.05, **P < 0.01, ***P < 0.001. **(G)** An example showing that gene co-expression module M4 and projection to CP-C share similar distributions on transcriptome UMAP (left). The correlation between M4 expression and CP-C projection intensity (right). **(H)** An example showing that gene co-expression modules M2 and projection to AId-I share similar distributions on transcriptome UMAP (left). The correlation between M2 expression and AId-I projection intensity (right).

Next, we analyzed whether the projection patterns of PFC neurons are interrelated to the co-expression of the neural circuit wiring molecules. We first analyzed the co-expression patterns of neural circuit wiring genes across all barcoded neurons, yielding five distinct co-expression modules (**Fig. 6E** and **S12B-D**). The relationship between projection patterns and gene co-expression modules was further analyzed, which revealed numerous significant correlations between the projection patterns and the strength of gene co-expression in specific modules (**Fig. 6F**). For example, the CP-C projection is highly related to the co-expression of M4 module, of which the projection intensity shows a positive correlation with the expression strength of M4 module genes (**Fig. 6G**). Similarly, the projection intensity targeting AId-I is significantly related to the expression strength of M2 module genes (**Fig. 6H**). Together, these results suggested that the neuronal projection patterns are closely related to the co-expression of molecules associated with cadherins and axon guidance.

### Predicting PFC neuron projection patterns using integrated transcriptomic and spatial information

The distinct expression patterns of neural circuits formation and maintenance related genes, such as cadherins and axon guidance genes in different projection classes inspired us to predict the projection pattern by gene profile via machine learning. While the gene profile can predict the projection pattern in certain degree [22, 29], the accuracy needs to be signicantly improved. Our data showed that on top of the correlation between neuron projection patterns with its gene profiles, the projection patterns are also highly related to its spatial location in the PFC (**Fig. 3G**). Thus, we tried to train a machine learning model to predict the projection patterns of PFC neurons based on transcriptome and spatial location. First, we integrated the single-cell transcriptomic dataset and spatial-omics dataset from SPIDER-Seq. This enabled us to obtain an integrated transcriptomic dataset with projection and spatial information at single-cell level. Then, we encoded binary labels (0 for no-projection or 1 for projection) for each targeted nucleus. Finally, we used the transcriptome principal components (PCs) and spatial information (spatial coordinates: X, Y, Z) as input features, and constructed an XGBoost machine learning model to predict the projection information (**Fig. 7A**).

**Figure 7.**
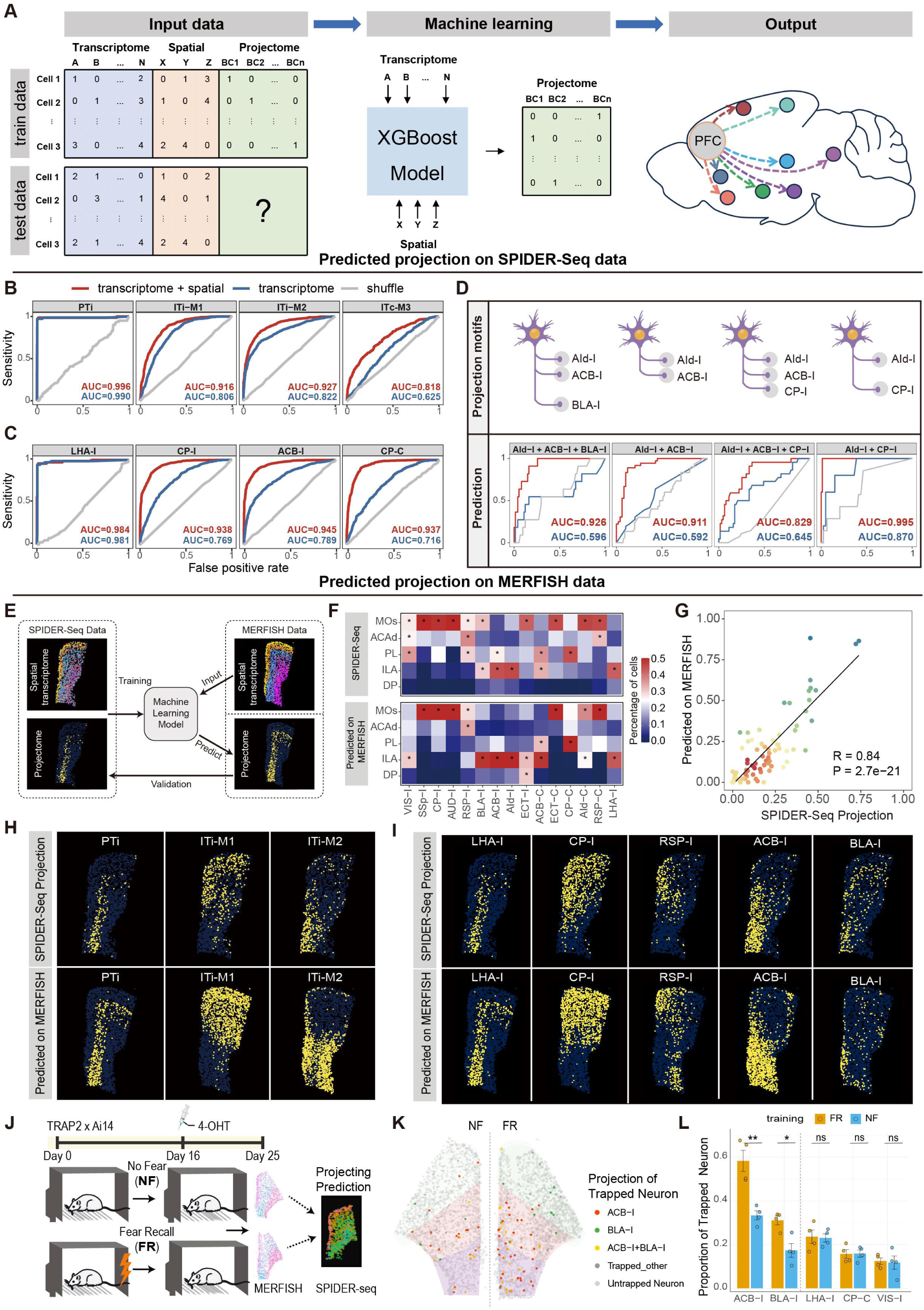
Prediction of neuron projection by integrated gene profiles and spatial location information by machine learning. **(A)** Schematic diagram of the steps in machine learning modeling. The train data (including transcriptome, spatial location, and projectome) was used to train an XGBoost model. The test data (including only transcriptome and spatial location) was used as the input of machine learning model and output the projectome information. **(B)** ROC curves for four predicted projection classes. The red curves unitize both transcriptomic and spatial profiles as input, the blue curves unitize only transcriptomic profile as input, and the gray curves are random shuffle control. **(C)** ROC curves for four predicted projection targets. The meaning of the curve color is the same as (B). **(D)** ROC curves for four predicted AId-I projection motifs. The meaning of the curve color is the same as (B). **(E)** Schematic diagram of predicted projection on MERFISH data. Our SPIDER-Seq data (including transcriptome, spatial location, and projectome) was used to train an XGBoost model. MERFISH data (including only transcriptome and spatial location) was used as the input of machine learning model and output the projectome information. Validation was performed by comparing the projectome predicted on MERFISH with the projectome mapped by SPIDER-Seq. **(F)** The percentage of neurons projecting to different nuclei in PFC subregions mapped by SPIDER-Seq (top). The percentage of neurons projecting to the putative projecting nuclei predicted by machine learning model based on the data from the corresponding MERFISH slice (bottom). Asterisks: percentage larger than 0.25. **(G)** Correlation between the predicted projections based on MERFISH data and the projections revealed by SPIDER-Seq. Each point represents the percentage of neurons in a PFC subregion for a target in (F). **(H)** The spatial distribution of neuron soma in different projection classes mapped by SPIDER-Seq (top), and predicted by machine learning model based on MERFISH data (bottom) (Bregma: 1.78mm). **(I)** The spatial distribution of neuron soma projecting to different targets mapped by SPIDER-Seq (top), and predicted by machine learning model based on MERFISH data (bottom) (Bregma: 1.78mm). **(J)** Re-illustration of experimental design from Sun et al.. TRAP2 × Ai14 mice were subjected to either fear recall (FR) or no-fear (NF) conditioning model. 4-hydroxytamoxifen (4-OHT) was administered on Day 16 to label behavior activated neurons. On Day 25, MERFISH spatial transcriptomics was performed on PFC. The MERFISH dataset were subjected to our machine learning model for projection prediction. **(K)** Representative MERFISH samples from NF (NF2_r0) and FR (FR2_r1) groups showing behavior activated (trapped) neurons. Predicted projection identities are color-coded: ACB-I (red), BLA-I (green), ACB-I+BLA-I (yellow) (Bregma:1.78mm). **(L)** Quantification of the proportion of trapped neurons within each major projection targets across four biological replicates. FR group shows significantly higher projections to ACB-I and BLA-I compared to NF (mean ± s.e.m.; *p < 0.05, **p < 0.01; two-sided t-test). Other projection targets show no significant differences (ns).

For model evaluation, we split our data into a training dataset (70%) and a test dataset (30%). The model was trained to predict both the overall projection classes (**Fig. 7B**) and specific projection targets for individual neuron (**Fig. 7C** and **S13A**), respectively. The receiver operating characteristic (ROC) curve demonstrated that our model achieved prediction performance with high accuracy. For example, the prediction accuracy of LHA-I projection pattern reached to 98.4%, and ACB-I with 94.5% accuracy. Notably, integrating spatial coordinates as input features significantly improved the model’s performance compared to using transcriptome features only (**Fig. 7B-C** and **S13A**). The lower accuracy of ITc-M3 may be due to its relatively less obvious transcriptomic and spatial features, while PTi shows significantly different transcriptomic and spatial features compared to IT neurons, which attributes to its higher prediction accuracy. To further demonstrate the accuracy of our model, we performed prediction at the level of projection motifs. Notably, our model can accurately predict projection motifs which are indistinguishable at the transcriptome level after integrating spatial information, such as the different projection motifs of AId-I (**Fig. 7D**).

To further validate the generalization capacity of our model, we tested it on the PFC MERFISH spatial transcriptome dataset from Bhattacherjee et al. [29]. First, we searched the MERFISH slice most closely matched with our PFC slice (**Fig. S13B**) and integrated both datasets based on the spatial and transcriptomic information (**Fig. S13C** and **D**). This allowed us to apply our projection model to the MERFISH data (**Fig. 7E**). The projection prediction on the MERFISH slices were highly consistent with the real projection experimental data revealed by SPIDER-Seq in the corresponding slice (**Fig. 7F**). The spatial visualization of the predicted results on MERFISH slice also shows consistent patterns with our SPIDER-Seq experimental data in the corresponding slice (**Fig. 7H-I** and **S13E**). The quantification analysis showed that the Pearson correlation between the overall predicted projection on MERFISH slice and the projection patterns on our slice resolved by SPIDER-Seq reached 0.84 (**Fig. 7G**).

Having verified the accuracy and generalization capability of our model in neural projection prediction, we next applied it to predict the projections of specific neurons under certain behavioral conditions, as a proof of concept. To achieve this, we obtained the MERFISH dataset from a prior study that captured fear recall (FR) engram neurons using the TRAP system. The experimental design of this study is illustrated in **Fig. 7J**. We aligned these MERFISH slices with our SPIDER-Seq data and integrated spatial and transcriptomic information to predict neuronal projections. Notably, we found that FR engram neurons projected significantly more to the ACB-I and BLA-I regions compared to non-fear (NF) control neurons. In contrast, no significant differences were observed in projections to other targets, such as the LHA, CP, and VIS (**Fig. 7K** and **L**). This finding aligns with previous reports implicating the invovement of PFC→ ACB-I and PFC→BLA-I pathways in fear recall-related behaviors [40–42].

## DISCUSSION

### Deciphering the multi-modal brain atlas by SPIDER-Seq

Neural circuit is an elaborating multi-modal network embedding diverse information such as connectivity, gene profiles, electrophysiology, and spatial location. Recent methods, such as, BARseq, BARseq2, VECTORseq, Retro-Seq, Epi-Retro-Seq, MERGE-Seq and Projection-seq have revolutionized high-throughput analysis of projectome information with some gene profiles [13, 17–19, 22]. However, there are still some urgent requirements for the improvements in: 1) reducing the cytotoxicity of tracer viruses which induces gene profile alterations; 2) detecting spatial location and the surrounding local environment information of the neurons in the network; 3) integrating multi-modal information; 4) simultaneous high multiplex targets tracing. To this end, we developed SPIDER-Seq by combining circuit tracing viral barcoding strategies with single-cell sequencing and spatial-omics, which allows us to integrate the transcriptome, projectome and spatial profiles of the neurons projecting to multi-nuclei with high throughput capacity and single-cell resolution. The integrity of SPIDER-Seq data was validated by the comparison between our data with previous fMOST [31], and spatial transcriptome data [29], as well as the data from different verifications [34]. This robust and cost-effective method can greatly facilitate to delineating the brain multi-modal network atlas and understanding neural circuit organization logic.

### The cellular and circuital spatial configuration and organization logic of the PFC

Although the molecular, cellular architecture and neural network of PFC have been extensively investigated, the integrated multi-modal PFC atlas embedding large scale neural circuit, transcriptomes, and spatial architecture information at single-cell resolution is still lacking, greatly hindering the understanding of the underlying cellular and circuital organization logic. By virtue of SPIDER-Seq, we delineated the detailed molecular, cellular and circuital spatial configuration of the PFC with single cell resolution. Consistent with previous studies, we showed that different transcriptomic cell clusters are arranged in different cortical layers with certain dorsal-ventral gradient, due to the inside-out development of these cell clusters in each layer [29, 43]. In contrast, the architectures of different projection motifs are configurated with a distinct geographical logic, in which each projection motifs gradiently occupy a specific territory in the 3D space of PFC. The neurons with the same projection pattern are allocated together in specific territory and consist of different transcriptomic cell clusters. By this organization principle, a single projection motif can more dynamically and distinctly decode neural signal inputs and produce more kinds of output signals by diverse expression of neurotransmitters/peptides and their receptors in their diverse transcriptomic cell clusters.

Although it has been demonstrated that majority of PFC IT projection neurons showed multiplex targeting and are sophisticatedly organized, the underlying organization logic remains elusive. Our SPIDER-Seq analysis revealed that the downstream nuclei targeted by the same projection class have significant higher probability to be co-projected by the PFC neurons, which may explain the overrepresented projection patterns which has also been observed by several studies [12, 44]. Moreover, the downstream nuclei targeted by the same PFC projection class have significantly more intensive reciprocal circuit connections. One possible underlying mechanism might be that the reciprocal connected neurons are more inclined to be both activated, and then simultaneously release certain kind of axon guidance cues. This might lead to the high probability to induce axon wiring from the same upstream nucleus during development. Functionally, the co-projection to the reciprocal connected neurons wiring principle may facilitate to information processing and the subsequent information exchange between these simultaneous signal recipients. It would be of great importance to examine whether this is a general logic for the whole sophisticate brain network.

### Specificity and complexity of neural signal decoding and transmission in the content of PFC neural network

Deciphering the dynamic and sophisticated neural signal transmission flow in neural network is the prerequisite for the understanding of neural computation and brain functions [35, 36]. However, the detailed signal decoding and transmission processes remain elusive. Owing to the high-quality single-cell transcriptome dataset integrated with projection information provided by SPIDER-Seq, we can detect the low expression genes such as receptor genes in content of neural network, which bring unprecedented insights into the logic of neural signal encoding in different circuits. In addition to the distinct neural signaling molecule expression patterns between PT and IT neurons, we also found significant differences in two projection classes of IT neurons, which may reflect the functional separation of PFC projections to the dorsal medial and the ventrolateral cortex as well as dorsal and ventral striatum [24, 34]. Notably, we showed that the expression pattern of diverse neural signaling molecules in different PFC circuits is not random organized; rather, it forms sophisticated and specific co-expression modules in different circuits, underlying the synergistic action among different neural signaling molecules.

The PFC expresses a variety of neurotransmitter receptor subtypes to decode the upstream neurotransmitter signaling, the different expression levels and combinations of receptor subtypes in PFC projection circuits may reflect the differential decoding of the same neurotransmitter input [37, 38]. In this line, we showed that individual PFC projection cluster selectively overexpress different receptor subtypes of the same neurotransmitter, underlying distinct neural signal decoding capacities and functions in different projection cluster. Moreover, for the projection neurons within the same projection motif, our SPIDER-Seq analysis revealed that they are gradiently distributed in the same spatial territory in the PFC and may likely receive the same local inputs. Thus, it would be important to distinctly encode the same inputs into more diverse signals via the combination of differentially expressed signal decoding receptor genes. By this principle, an identical input has the capacity to encode diverse kinds of downstream signals in the neural network, which may contribute to the generation of enough signal complexity to match the sophisticated information transmission capacity in the brain.

The release of neurotransmitters and neuropeptides is the primary means by which neurons transmit signals [39]. We found that different upstream projection neurons targeting the same nucleus showed distinct expression patterns of neurotransmitter/neuropeptide related genes, as illustrated by the distinct expression of neural neuropeptide genes in the IT projecting neurons targeting BLA from different upstream locations in the PFC. The molecular configuration of this kind of expression pattern may allow the same down-stream target nucleus to unambiguously differentiate inputs from different upstream nuclei, which might be important to guarantee the specificity of the neurotransmission in the intricate neural network. Together, our data depicted the landscape of neural information flow of PFC projection neurons and the underlying neural signal transmission molecules in this PFC network. Moreover, we discovered potential neural signal decoding and transmission principles to synchronize the specificity and complexity of neural transmission by sophisticated expression patterns of signal transmission molecules in different PFC projection neurons.

### Predicting PFC neuron projection patterns using integrated transcriptomic and spatial information

Our data revealed that different projection classes in the PFC distinctly express neural circuits formation and maintenance related genes, such as cadherins and axon guidance genes, which inspired us to predict the projection pattern by gene profile via machine learning. Consistent with previous studies [22, 29], while the gene profile can predict the projection pattern in certain degree, the accuracy needs to be further improved. As SPIDER-Seq can also decipher the spatial information of the projection neurons, which is also correlated to the projection pattern, we then integrated the spatial and the ample gene profile information at single-cell resolution for machine learning. By this approach, we achieved significantly high accuracy and, importantly, demonstrated generalization capacity of our model on the MERFISH dataset. Considering the projection neurons in other brain regions should, in principle, also have intrinsic correlations between the spatial gene profiling and projection pattern, our model can be likely used as an important basic foundation for fine-tune to predict projection patterns in any brain regions using a smaller amount of dataset. Thus, our data may further contribute to investigations such as, predicting neuron projection by spatial gene profiles after Ca2+ imaging or electrorheology experiments, which can integrate the function analysis with circuit information.

## LIMITATIONS

Despite the advances of SPIDER-Seq, there are several issues that need to be further improved: 1) Viral labelling coverage: This is limited by the diffusion area in the injection sites, especially for targeting large nuclei such as the CP. To tackle this issue, we performed injections at different depths to increase the coverage area of the virus (see Methods); 2) Viral infection efficacy: Our previous work reported that rAAV2-retro has high efficiency for most cortical area labeling, but relatively lower for other regions [45]. To increase the labelling efficacy, we waited for 30 days to reach high infection efficacy. In future, it would be important to engineer new generation of highly efficient tracing virus; 3) Local circuit mapping: SIPDER-Seq applied for local circuit mapping, such as inhibitory neuron circuits, is subjected to future study. 4) It should be noted that our study used only females, but compared with male datasets [29, 31]. Although, there are no reports indicating differences in PFC circuit connectivity between female and male mice, this will remains an important consideration for future research on sex-related study.

With the rapid advancing research in interdisciplinary fields, such as viral bioengineering, in situ sequencing methods, imaging technologies, and machine automation, SIPDER-Seq can be further upgraded to delineate the integrated spatial transcriptome and projectome atlas with high throughput, high 3D resolution, more accuracy, multi-modal information. Moreover, the integrated multi-modal neural network will also shed insights into the organization logic and the computation principle of neural network and ultimately contribute to the brain-inspired artificial intelligence.

## MATERIALS AND METHODS

### Animal care

Animal care procedures and experiments (HZAUMO-2024-0318) were approved by the Scientific Ethics Committee of Huazhong Agricultural University, Hubei, China, and conducted ethically according to the Guide for the Care and Use of Laboratory Animals of the Research Ethics Committee of Huazhong Agricultural University. Mice were housed in facility with a standard light cycle (12 hr light/12 hr dark) and ad libitum access to food and water.

### rAAV2-retro virus Barcoding

The rAAV2-retro barcode core plasmid (Plasmid #32395) was generated by replacing CMV-GFP-poly(A) with hSyn(452 nt)-GFP-BGH (225 nt), followed by insertion of NheI and BglII sequence downstream of the GFP. All barcode sequences were inserted between NheI and BglII by enzyme ligation or homologous recombination. We constructed 33 core plasmids carrying different barcode sequence (**Table S1**). The barcoding virus was rescued by transfecting the rAAV2-retro barcode plasmid, pAdDeltaF6 (Plasmid,112867) and rAAV2-retro helper plasmid (Plasmid,81070) into 293T cell line (ATCC,CRL-3216). The final product was purified by ioxanol ultracentrifugation to obtain the SPIDER-Seq tracing virus. The virus titer was adjusted to 2×10^12^ viral particles per mL.

### Virus Injections

Female C57BL/6 mice at 7-8 weeks age were anaesthetized with a mixture of anesthetics 65 mg/kg ketamine and 13 mg/kg xylazine (i.p. injection). Their heads were fixed to a stereotaxic apparatus (68030, RWD, China). After exposed the skull, holes on the skull surface were drilled with corresponding coordinates (AP axis and ML axis, the coordinates for each nucleus are shown in **Table S2**). The virus was injected at a rate of 40 nl/μl, and 150 nl per nucleus. The micropipette was left in the tissue for 5 min before and after injection, to prevent virus spilling and backflow. The above operations were repeated to complete the injection of all nuclei (12-16 nucleus per mouse). The entire operation lasted about 6-8 h with 50 μl of anesthetic replenished every 3 hours. To compared the barcode in situ sequencing results with the viral tracing results of rAAV2-retro expressing fluorescent protein (EGFP or mCherry), we injected 150 nl of the rAAV2-retro expressing fluorescent protein into one target nucleus in each mouse. After surgery, the mice were placed on a heating pad to allow recovery and were allow housed under a 12-hour light/dark cycle at 22-25 °C for one month.

### Single cell dissociation

One month after barcode virus injection, mice were anesthetized with isoflurane and decapitated. The brain was immediately removed and then sectioned into 300 μm slices in ice-cold ACSF (124 mM NaCl, 2.5 mM KCl, 1.2 mM NaH2PO4, 24 mM NaHCO3, 5 mM HEPES, 13 mM glucose, 2 mM MgSO4, and 2 mM CaCl2, pH: 7.3-7.4) on vibratome (Leica VT1200). Slices containing PFC region were transferred into Petri dish containing ice-cold ACSF with 45 µM Actinomycin D (Sigma-Aldrich, Cat# A1410). PFC region containing the anterior cingulate area (ACA), prelimbic area (PL), infralimbic area (ILA), secondary motor cortex (MOs), medial orbital cortex (ORBm), Dorsal peduncular area (DP) (∼Bregma 2.58 mm - Bregma 0.86 mm according to Paxinos and Franklins the Mouse Brain in Stereotaxic Coordinates) labelled with green fluorescence were isolated under a fluorescence microscope. Single cell suspensions were prepared as described previously [46]. The isolated tissues were quickly cut into small pieces less than 1 mm and transferred to digestion buffer containing 3 mg protease XXIII (Sigma-Aldrich, P5380) and 30 U/ml papain (Sigma-Aldrich, P3125). The digestion was performed at 34°C for 30 min and bubbled with a mixturegas of 95% O2 and 5% CO2 continuously. After the digestion, the tissue was transferred to stop buffer (ACSF contain 1 mg/ml Trypsin Inhibitor (Sigma-Aldrich, T6522), 2 mg/ml BSA (Sigma-Aldrich, A2153) and 1 mg/ml Ovomucoid Protease Inhibitor (Worthington, LK003153). We gently titrated the digested tissue with 4 polished Pasteur pipets, of which the bore diameter of the pipets is successively decreasing from 600 µm to 150 µm. Following trituration, suspension was filtered through a 30 mm filter, then centrifuged at 300 g for 5 minutes, The pellet was then resuspended in 700 µL ice-cold, carbogen-bubbled ACSF with 0.01% BSA and then subject to single cell RNA sequencing library preparation using BD Rhapsody single-cell Analysis System (BD Biosciences, 633702) according to the manufactory’s manual.

### Single-cell RNA sequencing library preparation

Single-cell mRNA sequencing library was performed by the BD Rhapsody Single-Cell Analysis System (BD Biosciences, 633702). Firstly, single-cell suspension in BSA was loaded into a BD Rhapsody cartridge (BD Biosciences, 633733) with >200,000 microwells. Secondly, single-cell mRNA was captured by magnetic beads with barcoded capture oligos (BD Biosciences, 664887). Then, magnetic beads were collected for cDNA synthesis and library construction following the BD Rhapsody single cell 3’ whole transcriptome amplification (BD Biosciences, 633733) workflow. Finally, the libraries were sequenced on Illumina NovaSeq 6000 (Illumina, USA) with 300-bp reads (150-bp paired-end reads).

### *In situ* Sequencing of the transcripts of PFC genes and barcodes

Probe design: We modified the protocol according to previous in situ sequencing method [32]. Briefly, 4-20 pairs of probes were designed for each gene depending on the length of mRNA (marker gene or barcode), and all probe sequences were shown in **Table S3**. Each end of the padlock probe contains 13 nt complementary to the target sequence, and the middle region contains two repeats of the complementary sequence of the detection probe. The 5’ and 3’ end of initiator primer are complementary to the 3’ end of the targeting sequence and padlock probe, respectively. We customized 12 groups of 16 bp fluorophore-modified detection probes (Alexa Fluor 488, CY3, CY5, and CY7) (**Table S4**). 47 gene are detected by 12 imaging cycle and five channels for DAPI, 488, cy3, cy5, cy7 are used in each cycle to rearch more robust and efficient results. The padlock probe was phosphorylated with T4 Polynucleotide Kinase at 200 µM (Vazyme, N102-01), and then annealed with initiator primer. Detection probes were dissolved at 100 µM in ultrapure RNase-free water. All probes were stored at -80°C before use.

Slices preparation: One month after virus injection, the mice were perfused with 4% PFA, and postfixed in 4% PFA for 24 h. We then cryoprotected the brain with 30% sucrose until tissue sinks and embed the mice brain in OCT. The brain tissue was sliced to 15 µm continuously with a Leica cryostat (Leica CM3050 S). Slices containing PFC were mounted on the poly-L-lysine pretreated cover glass then stored at -80°C until used. The brain slices were sealed in Secure-Seal hybridization chambers (Grace, 621505), fixed with 4% PFA for 10 mins, permeabilized with pre-cooled methanol at -80°C for 15min, then followed by pepsin (2 mg/mL, Sigma, P0525000) digestion at 37°C for 90 s.

The probes were diluted at 25 µM to 100 µM in hybridization buffer with 2X SSC (Sangon, B548109), 10% formamide (Sangon, A100606) and 20 mM RVC (Beyotime, R0108)) and incubated overnight at 37°C. The brain slices were washed twice in PBSTR (0.1% Tween-20, 0.1 U/µL RRI in PBS) for 20 min, followed by wash in 4X SSC dissolved in PBSTR. Next, the brain slices were incubated in the ligase reaction mixture (1 U/µL SplintR ligase (NEB, M0375L), 1X buffer, 0.2U/µL RRI) for 2 hours at 25°C. Following 20 mins wash with PBSTR, and then incubated with RCA mixture (1 U/µL Phi29 (Vazyme, N106-01), 1X RCA buffer, 0.25 µM dNTP, 0.2 µg/µL BSA, 5% Glyceryl)) at 30°C for 6 hours.

Imaging: Slices were incubated with detection probe labelled with fluorescence (488, cy3, cy5, cy7) at 37°C for 30min in each round, and washed with PBST for 3 times, following with DAPI staining. Slice were scanned by Leica THUNDER Imager system with 20× lens (NA 0.75). After each round of imaging, signals were stripped by stripping buffer (60% formamide in 2X SSC) twice at room temperature for 10 mins for each round. DAPI staining was performed 3 cycle of image.

### scRNA-seq and barcode data pre-processing

Raw reads were pre-processed using the BD Rhapsody™ Whole Transcriptome Analysis (WTA) pipeline (v1.11) (https://bd-rhapsody-bioinfo-docs.genomics.bd.com). The R1 reads were analyzed to identify the cell label sequences (CLS), common linker sequences (L), and Unique Molecular Identifier (UMI) sequence. The R2 reads were used for aligning to the reference genome and annotating genes. For WTA reference genome, we selected GRCm38-PhiX-gencodevM19-20181206.tar file. For transcriptome annotation, we selected gencodevM19-20181206.gtf file. After setting up, we ran pipeline using the default parameters. The expression matrix file generated by pipeline was used for transcriptome analysis, while the BAM file was used for extracting projectome information.

### scRNA-seq quality control

Single cell RNA-seq transcriptome analysis is mainly performed by R package Seurat (v4.4.0) [47]. Briefly, Seurat object was created using the “CreateSeuratObject” function, and the gene expression profile of each cell was then normalized using the “NormalizeData” function with scale.factor = 10000.

We filtered the following cells: nCount_RNA < 1000, nFeature_RNA < 1000, and mitochondrial contents > 15%. We removed neuronal cells with nFeature_RNA < 1500, because previous studies showed that neuronal cells have a higher number of genes expressed than non-neuronal cells [27]. The following genes were filtered: min.cells < 3, mitochondrial genes, and ribosomal genes. Then, the R package DoubletFinder (v2.0.3) [48] was used to remove potential doublets.

### Clustering of scRNA-seq transcriptome

First Seurat Canonical Correlation Analysis (CCA) was used to remove batch effects between three samples. The “SelectIntegrationFeatures” function was applied to select top 2000 common variable genes across three samples. Then we used the “FindIntegrationAnchors” function to find anchors, and integrated the three datasets together using the “IntegrateData” function. After integration, we performed the standard Seurat clustering analysis workflow. We used the “ScaleData” function to scale the integrated data, and performed principal component analysis (PCA) using the “RunPCA” function. Then we computed the nearest neighbors used the “FindNeighbors” function with top 20 PCs. We used the “FindClusters” function for clustering analysis with resolution = 0.5. Then, we annotated the clusters based on previously reported markers of PFC cell types [29]. We manually removed some mixed low-quality cell clusters that expressed markers of multiple cell types. Seven main cell types were annotated: Excitatory neuron, Inhibitory neuron, Astro, Endo, Microglia, Oligo, OPC. Then, we extracted Excitatory cell type for further analysis. We used the “FindClusters” function for Excitatory clustering analysis with resolution = 2. Then, we annotated the clusters based on previously reported markers of PFC Excitatory subtypes. We annotated a total of 13 Excitatory subtypes: L2/3_IT_1, L2/3_IT_2, L4/5_IT_1, L4/5_IT_2, L5_IT_1, L5_IT_2, L6_IT_1, L6_IT_2, L5_PT_1, L5_PT_2. Then, we ran the Uniform Manifold Approximation and Projection (UMAP) dimensional reduction using the “RunUMAP” functions, and visualized data using functions provided by Seurat. In total, our data contains a transcriptome expression matrix of 33,766 cells and 26,902 genes. 9,038 neurons were labeled with barcodes (1,842 for mouse1; 4,410 for mouse2; 2,786 for mouse3).

### Projectome barcode alignment

We constructed a ‘database.fasta’ file containing our barcode sequences, and used the blast makeblastdb command (v2.13.0) to produce BLAST databases. Then, we used the samtools (v1.13) [49] software to extract R2 reads sequences, cell labels, and Unique Molecular Identifier (UMI) sequences from the BAM file, and constructed a ‘query.fasta’ file. We used the blastn command to align the ‘query.fasta’ file with the barcode BLAST database using the following parameters: -task blastn-short -word_size 4 -evalue 1 -outfmt “6 qseqid sseqid nident” -max_hsps 1 -ungapped -num_threads 10 -mt_mode 1. We kept sequences with 3 or fewer mismatches and removed sequences with duplicate UMI to generate the cell-barcode projectome expression matrix.

### Removing barcodes background noise

We observed low expression of barcodes in some non-neuronal cells in the projectome expression matrix. Due to the slightly different sequencing depths of the 3 scRNAseq samples, we set different thresholds for different samples to filter out background noise accordingly. First, we divided the cells into Neuron and Non-neuron categories and plotted the UMI Counts-Barcode curve. We used the UMI Counts of the elbow point position of the Non-neuron curve as the threshold, and set the barcode expression below the threshold in Neuron and Non-neuron as 0. Finally, to compare the projection patterns between neurons on the same scale, we performed Min-Max Normalization on the expression levels of barcodes in each cell. With that, the raw barcode counts were rescaled between zero and one.

### Clustering of single cell projectome

R stats (v4.2.0) “hclust” function was used to perform hierarchical cluster analysis for the cell-barcode matrix. The “cutree” function was then applied to cut the tree into clusters at a height=1.5. After manually merged clusters with similar barcode expression profiles, we obtained a total of 33 projectome clusters. Then, we run UMAP dimensional reduction on the cell-barcode matrix, and merged 33 projectome clusters into 4 projectome modules.

### Visualization of the distribution of projection neurons on UMAP

Density scatter plots were used to visualize the distribution of different projection neurons on the UMAP. Specifically, we used the “geom_pointdensity” function from R package ggpointdensity (v0.1.0) to create scatterplots where each point was colored by the number of neighboring points. This is useful to visualize the 2D-distribution of points in case of overplotting.

### Calculation of projectome correlation between SPIDER-Seq and fMOST data

To verify the reproducibility of SPIDER-Seq projectome data, we collected fMOST projectome data from Gao et al. [31]. It contains the number of neurons that PFC project to downstream targets (**Tables S6**). We compared the number of neurons project to each target with our SPIDER-Seq data. We plotted the correlation curves and calculated the Pearson correlation coefficient (**Fig. S1O**). Each point represents a target, the X and Y axes represent the projection intensity in fMOST and SPIDER-seq (Min-Max normalization). The correlation between SPIDER-seq and fMOST projections reached 0.81.

### Calculation of projection motifs

To calculate the statistically significant projection motifs, we constructed a null model based on a previous study [12]. Briefly, we first assumed that each neuron projected to each target were independent, and used the binomial cumulative distribution function to calculate the expected cell number of each projection motifs. We then calculated the P value by comparing the expected cell number to the observed cell number, and corrected the P value with the Bonferroni method. We defined the projection motifs with log2 fold change > 1 or < -1 and P value < 0.01 as significantly over- or underrepresented projection motifs.

### Calculation of the correlation between projectome, spatial location, and transcriptome

The spatial, transcriptome, and projectome distances for different projection motifs were calculated to plot the correlation scatter plot. Specifically, we used the “dist” function in R to calculate the euclidean distance between each pair of projection motifs. The “geom_pointdensity” function from R package ggpointdensity (v0.1.0) was used to create the scatterplots. Correlation was calculated using the Pearson correlation coefficient.

### Image registration and cell segmentation

The raw images were registered for each round using BigWarp (v9.1.2). Specifically, we used the first round image as the reference and manually aligned the corresponding manually labelled“aligning points” between each round image with the first round image in BigWarp’s “landmark mode”. After alignment, we performed image registration. We used FIJI to stack all channels and rounds of images of the same slice into one image file. We used the DAPI channel of each slice to perform cell segmentation by Cellpose (v2.0.5) [50] and then calculated the average gray value of each channel in each cell to generate the spatial cell-channel expression matrix.

### Spatial expression matrix quality control

We manually set thresholds for quality control on the spatial cell-channel expression matrix to filter out values with low channel expression to remove background noise. We filtered out cells with a volume smaller than 50 pixels (32.5um^2^) or larger than 500 pixels (325um^2^) in each slice, as well as the cells with total expression levels less than 5 or higher than 400, to remove low-quality cells and potential doublets. Then, we loaded the expression matrix into Seurat and used the “NormalizeData” function for normalization, with the parameter scale.factor=100.

### Spatial cell annotation

To annotate cells in the spatial data, we used Tangram (v1.0.4) [33] to map the annotated information of scRNAseq clusters onto the cells in the spatial-omics map. Briefly, we used “tg.pp_adatas” function to find the common genes between adata_sc and adata_sp. Then we used the “tg.map_cells_to_space” function to perform the cluster level mapping, with parameters: mode=’clusters’, cluster_label=‘SubType’.

### PFC 3D visualization

For 3D visualization of PFC, we used R package WholeBrain (v0.1.1) [51] and extended functions [29] to align the spatial slices with Allen Brain Atlas CCF v3. First, each brain slice was paired to the closest matching coronal section in CCF v3 with the help of DAPI image and spatial location of the cell types. Then, we manually adjusted the scale and position of each brain slice to ensure accurate alignment with CCF v3. After alignment, we extracted the PFC regions according to the brain region annotation information in CCF v3. Then, we used R package rgl (v1.1.3) and Allen brain 3D mesh [43] to visualize PFC in 3D.

### Single cell weighted gene co-expression network analysis

Single cell weighted gene co-expression network analysis is mainly performed by R package hdWGCNA (v0.3.01) [52]. Briefly, “SetupForWGCNA” function was used to select neural signal molecule or neural circuit wiring molecule genes for analysis. We used “MetacellsByGroups” function to construct metacell expression matrix, and normalized the matrix using “NormalizeMetacells” function. Then, “SetDatExpr” function was used to specify the expression matrix for network analysis, and soft power threshold was selected using “TestSoftPowers” function. We used “ConstructNetwork” function to construct the co-expression network, and visualized the network using functions provided by hdWGCNA. Module Eigengenes (MEs) are a commonly used metric to summarize the gene expression profile of an entire co-expression module. Briefly, module eigengenes are computed by performing principal component analysis (PCA) on the subset of the gene expression matrix comprising each module. The first PC of each of these PCA matrices are the MEs. We used the “ModuleEigengenes” function from R package hdWGCNA (v0.3.0.1) to calculate the MEs for each gene co-expression module. To fit the correlation curve, we divided cells into 10 bins, ranging from 10% to 100%, based on the MEs values. We then calculated the average projection intensity and MEs for each bin and plotted the correlation curve.

### Neuron projection prediction by machine learning

We trained a XGBoost model to predict projectome by transcriptome and spatial location in each neuron. In order to simultaneously obtain the whole transcriptome and spatial location of single neuron, we calculated the projectome correlation coefficient between each cell in scRNAseq dataset and spatial-omics dataset according to their embedded projection information in both datasets. Based on the projection similarity, the spatial location (X, Y, Z) of a neuron in the spatial-omics dataset was assigned to the neuron in scRNAseq dataset with the highest projection correlation coefficient. We used the R package caret (v6.0-94) “ createDataPartition” function to split the scRNAseq cells into training and test datasets with a 7:3 ratio. The first 30 PCs of the transcriptome and the spatial X, Y, and Z coordinates were used as features, and the binary annotation of the projection module or target area were used as the labels. We first used the R package xgboost (v1.7.5.1) “xgb.cv” function to find the optimal nround, and then used the “xgboost” function to train model, using the following parameters: max_depth=5, eta=0.5, nthread = 5, nround = xgb.cv$best_iteration, objective = “binary:logistic”. Next, we used the “predict” function to make predictions on the test dataset. ROC curve and area under the curve (AUC) values were obtained using R package PRROC (v1.3.1) “roc.curve” function. To further verify the generalization ability of our XGBoost model, we performed projectome predictions based on PFC MERFISH data from previous a study [29]. We first integrated the MERFISH data with SPIDER-Seq data using Seurat’s standard CCA analysis pipeline. Then the first 30 PCs of the transcriptome and the X and Y coordinates were used as features. SPIDER-Seq data were used as the training data and the PFC MERFISH data were used as the test data.

## DATA AVAILABILITY

The raw single cell RNA-seq data are available from GEO (<u>GSE273066</u>). The raw spatial-omics data for this study are available via Hugging Face at https://huggingface.co/TigerZheng/SPIDER-STdata. The processed data ready for exploration can be accessed and downloaded via our interactive browsers at https://huggingface.co/spaces/TigerZheng/SPIDER-web (**Fig. S14**).

## CODE AVAILABILITY

All data were analyzed with standard programs and packages. The codes were freely accessible from https://github.com/ZhengTiger/SPIDER-Seq.

## ACKNOWLEDGEMENTS

We thank the experimental platform of National Key Laboratory of Agricultural Microbiology of Huazhong Agricultural University. We thank Spatial FISH, Co., Ltd for assistance with single-cell RNA sequencing and spatial multi-omics.

## AUTHOR CONTRIBUTIONS

G.C., LQ.S., JX.D. conceived and designed the project. LQ.S., Y.W., GH.D. designed the barcoded virus. LQ.S., YY.H. constructed and purified the barcoded virus. LQ.S. performed multiple injections into the mouse brain. LQ.S., ZC.W., LY.Y., YY.H. contributed single cell dissociation. KJ.Y. prepare single-cell RNA-Seq libraries. LQ.S., XH.H., XF.W., YY.H. performed fluorescence in situ hybridization. H.Z. performed SPIDER-Seq data processing pipeline development. H.Z., YP.Y., XJ.G. performed single cell sequencing data analysis. H.Z., YY.H. performed spatial omics data analysis. H.Z. performed SPIDER-web online website development. G.C., LQ.S., H.Z. wrote the manuscript. YY.H., DQ.M., H.C., and HZ.L. revise the manuscript. All authors read and approved the final paper.

---

**Figure S1.**
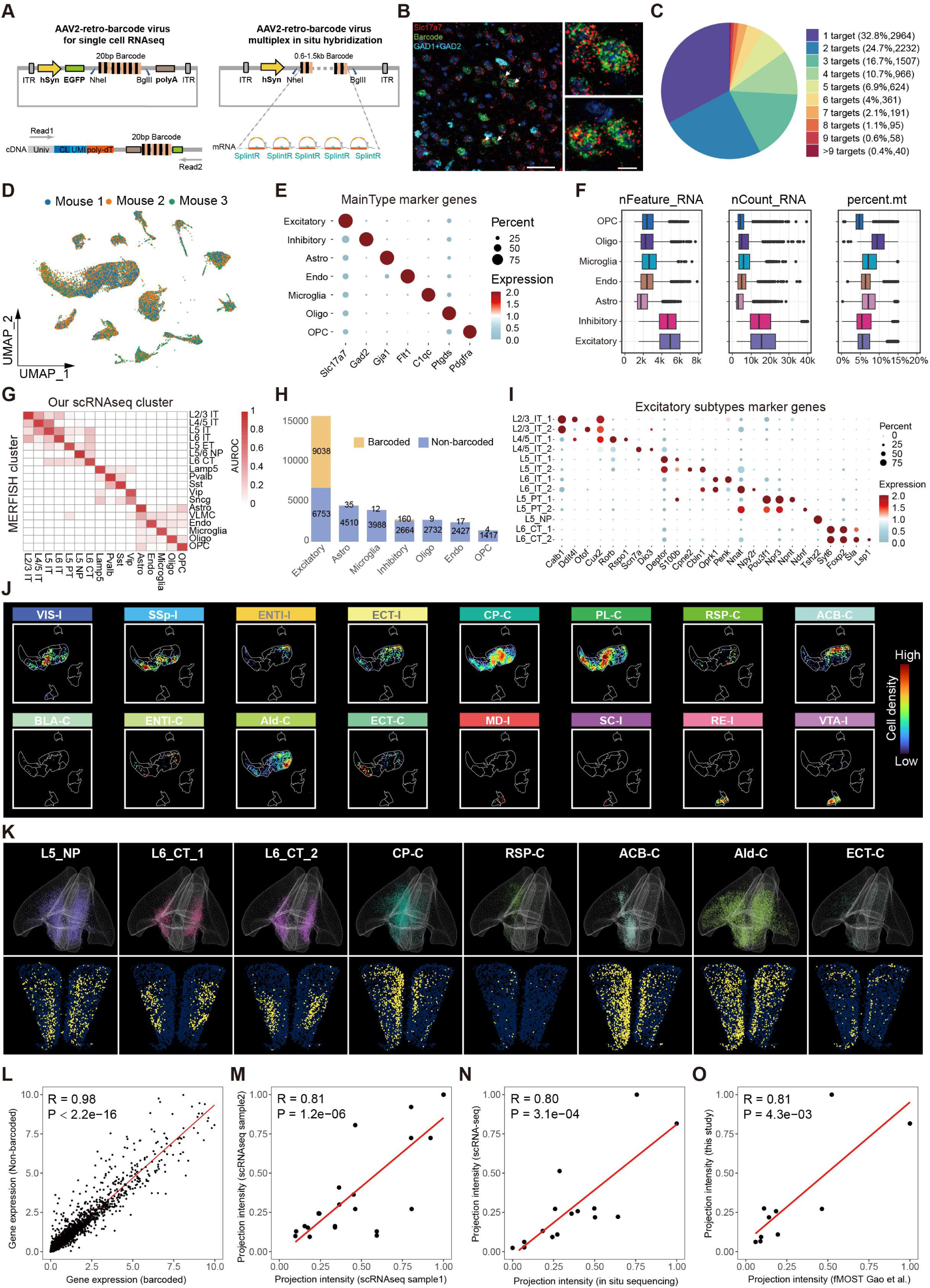
Delineating multi-modal PFC atlas embedding single-cell projectomics, spatial-omics and transcriptomics by SPIDER-Seq. **(A)** rAAV2-retro-barcode virus designed for single cell RNAseq and schematic of designed primers to recover cell barcode and UMI in read 1, and 3’ tail of EGFP and virus barcode in read 2 (left). rAAV2-retro-barcode virus designed for multiplex in situ hybridization and schematic of designed hybridization primers to detect barcode (right). **(B)** Fluorescence in situ hybridization detection shows that the retrograde tracing barcode was mainly distributed in excitatory neurons. Excitatory neuron marker: Slc17a7, red; Inhibitory neuron markers: Gad1 and Gad2, blue; Barcode signal retrograde tracing from ACB: green. Scale bars, 50 µm. The inset on the right shows a magnified view of the overlap of Slc17a7 and ACB barcode. Scale bars, 10 µm. **(C)** Distribution of number of projection targets of PFC barcoded neurons. **(D)** Integrated UMAP of cells from 3 mouse brains (mouse1:12 targets, mouse2: 14 targets, mouse3: 16 targets (Table S2)), colored by samples. **(E)** Dotplot showing the expression patterns of maintype marker genes in transcriptome maintypes. **(F)** Boxplots showing the distribution of the number of genes (left), number of UMI (middle) and mitochondrial genes percentage (right) detected in each transcriptome maintype. **(G)** Heatmap showing the gene expression correlation between the PFC clusters defined by scRNA-seq in SPIDER-Seq and Bhattacherjee et al.. **(H)** Barplot showing the number of barcoded (yellow) cells in each transcriptome maintype. **(I)** Dotplot showing the expression patterns of excitatory marker genes in excitatory subtypes. **(J)** Density scatter visualization on UMAP of PFC neurons projecting to different nuclei. **(K)** Spatial distribution of different transcriptome subtypes and projection neurons to different nuclei in 3D (top) and a 2D example slice (bottom) (Bregma: 2.1mm). **(L)** Correlation of transcriptome in barcoded and non-barcoded neurons. **(M)** Correlation of projectome in different scRNAseq samples. **(N)** Correlation of projectome in scRNA-seq and spatial-omics data. **(O)** Correlation of projectome revealed by our SPIDER-Seq and fMOST data.

**Figure S2.**
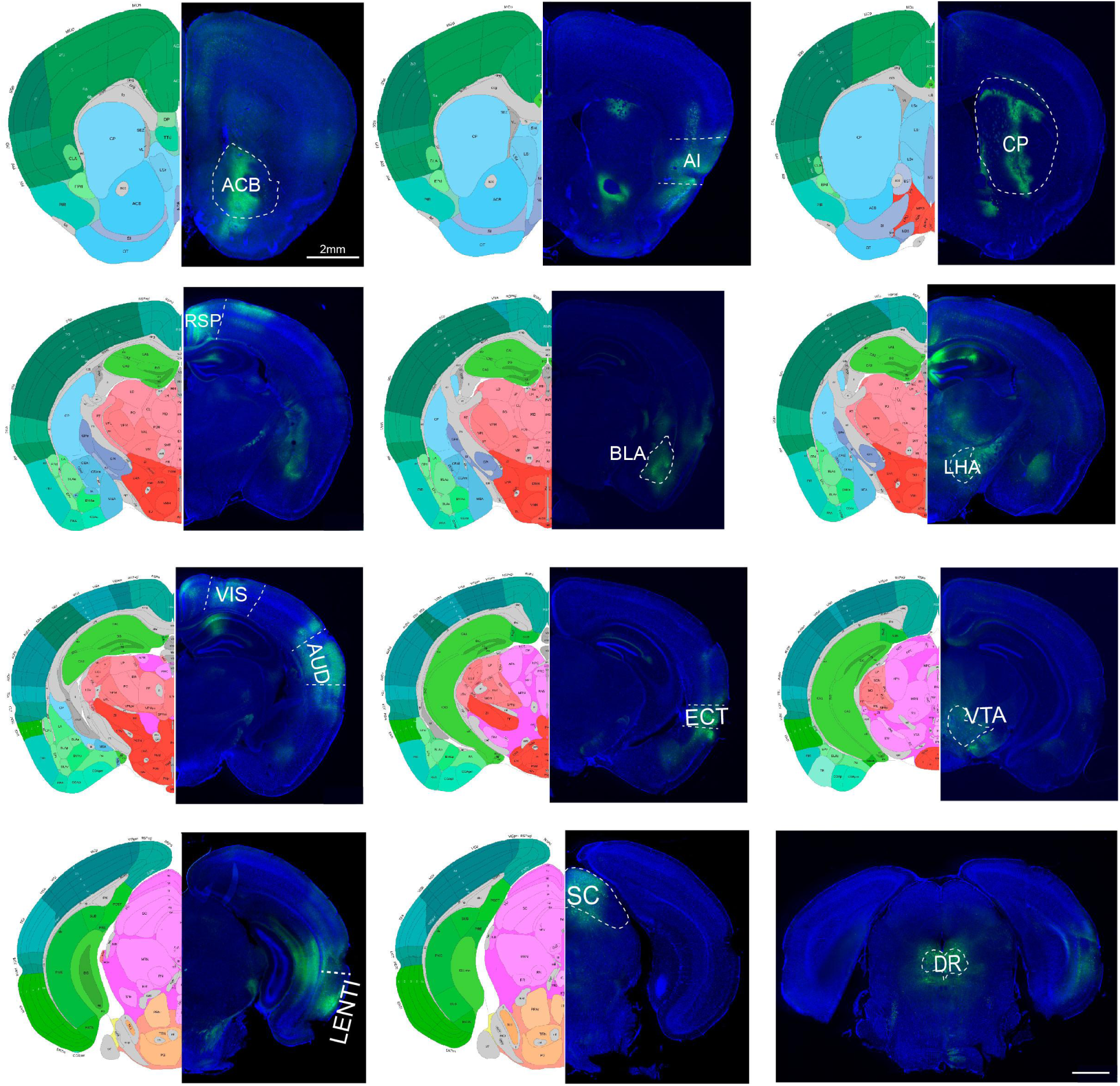
Injection sites of different nuclei targeted by PFC. Coronal images of the injection sites (ACB, AI, CP, RSP, MD, BLA, LHA, VIS, AUD, ECT, VTA, LENTl, SC, DR) of rAAV2-retro-barcode virus (right) and the corresponding Allen brain atlas (left). Scale bar: 2 mm.

**Figure S3.**
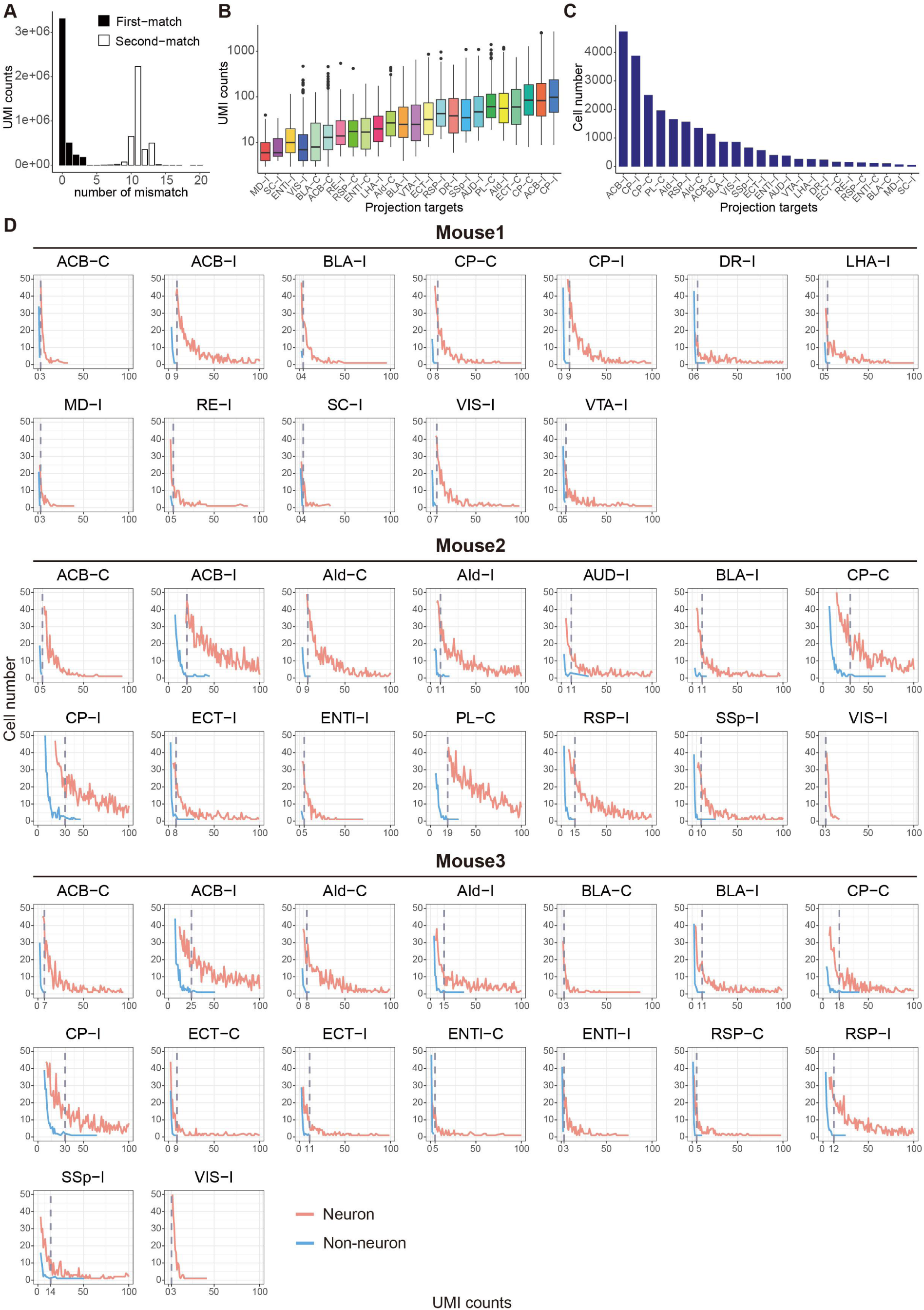
Extracting projection information from scRNAseq. **(A)** Histogram of the number of mismatches between the First-match reads and the Second-match reads. This indicates that there is enough sequence diversity to distinguish the correct barcodes. **(B)** Box plot shows the UMI counts for the barcode projecting to different nuclei. **(C)** Histogram shows the number of barcoded cells for 24 targeted nuclei. **(D)** The UMI counts and cell number curves for each barcode in 3 scRNAseq samples, grouped by Neuron and Non-neuron. The gray dashed line indicates the threshold of UMI counts for a barcode filter after elbow analysis.

**Figure S4.**
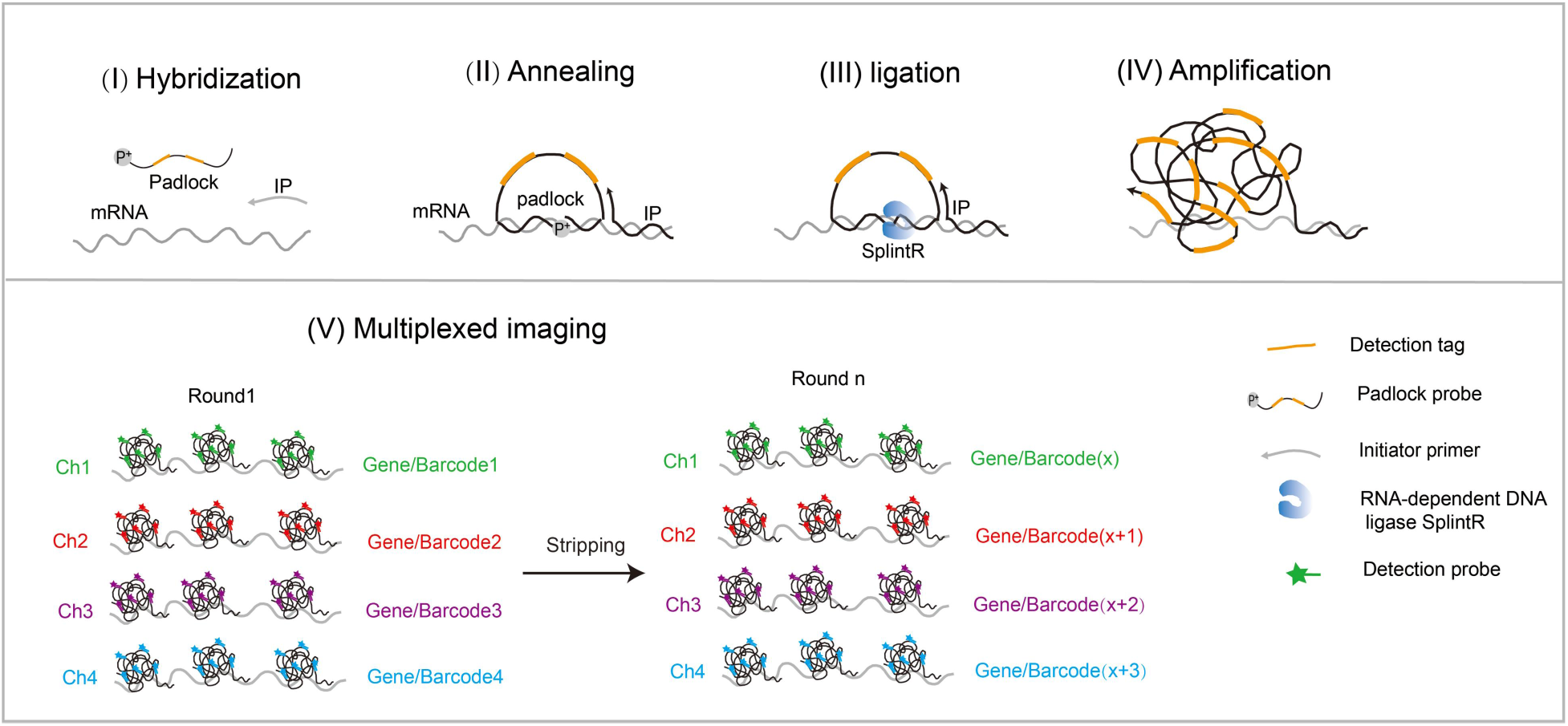
Flow diagram of multiplex detection based on a modified version of MiP-seq. I. The padlock probe and initiator primer (IP) are hybridized to the target RNA. Notably, the dual barcode primer in MiP-seq is replaced by two repeating detection tag. II. Padlock probe and IP are annealed to the target mRNA to form a padlock structure. III. Padlock probe is circularized by RNA-dependent DNA ligase SplintR to form the RCA template. IV. RCA of circled padlock probe forms rolling-circle products. V. The detection tag is hybridized to the detection probe labelled with fluorescence. After each imaging cycle, probes are stripped away from the tissue using 60% formamide for the next cycle. Five channels for DAPI, 488, cy3, cy5, cy7 are used in each round and each channel represents a detected gene. We detected 47 genes in 12 rounds (**Table S4**).

**Figure S5.**
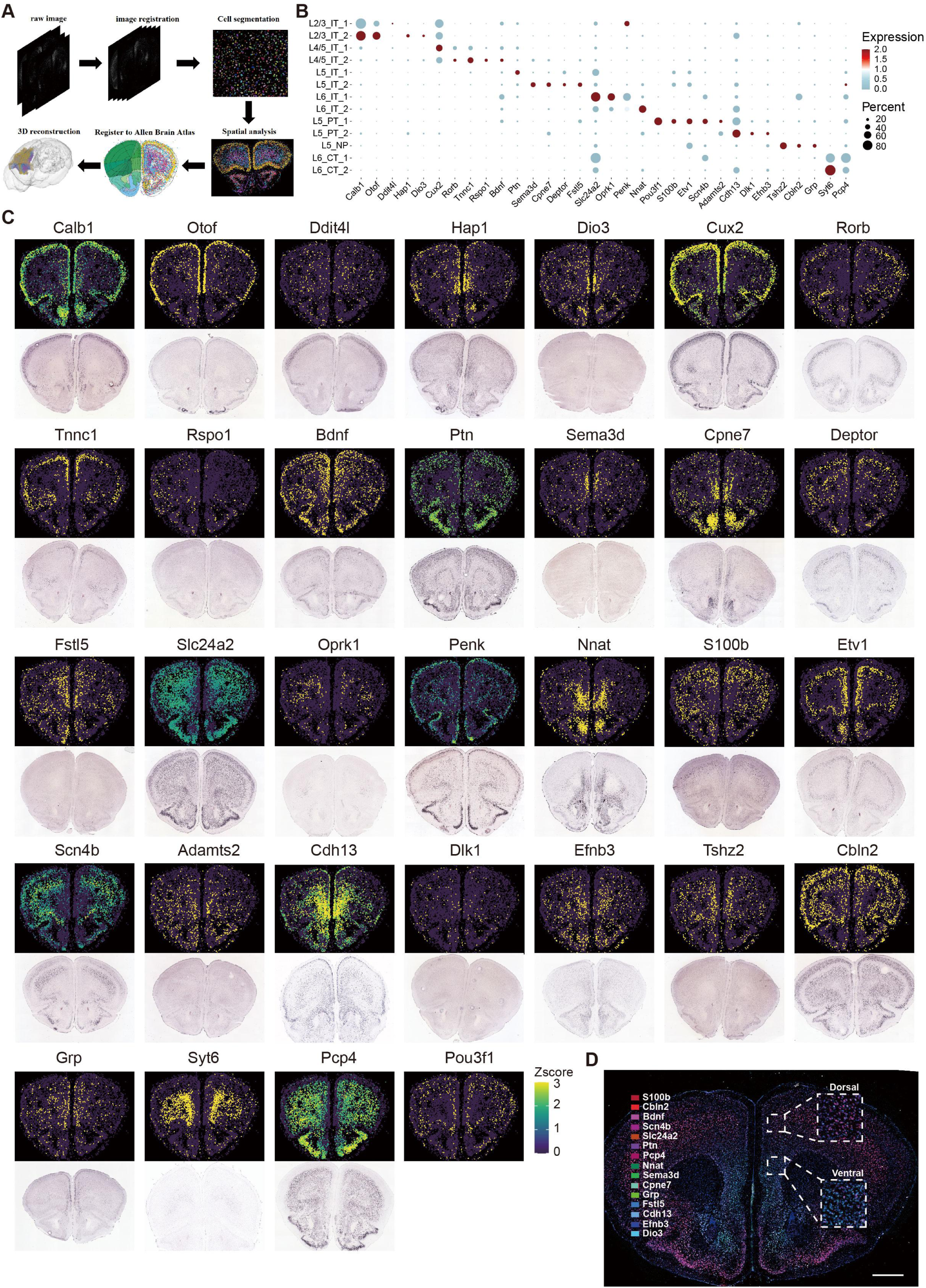
Spatial gene profiles data analysis of SPIDER-Seq. **(A)** Spatial expression data analysis process of SPIDER-Seq, including image registration, cell segmentation, spatial analysis, registration to Allen brain atlas, and 3D reconstruction. **(B)** Dotplot showing the expression patterns in excitatory neurons subtypes of 32 marker genes measured by SPIDER-Seq. **(C)** In situ hybridization images show that the expression pattern of 32 markers detected by SPIDER-Seq (top) are consistent with ISH data from Allen Brain Atlas (bottom) (Bregma:2.1mm). **(D)** The spatial transcription shows spatial gradient separation of the PFC transcription atlas with dorsal-enriched genes (*S100b*, *Cbln2*, *Bdnf*, *Scn4b*, *Slc24a2*, *Ptn*, *Pcp4*) and ventral-enriched genes (*Nnat*, *Sema3d*, *Cpne7*, *Grp*, *Fstl5*, *Cdh13*, *Efnb3*, *Dio3*).

**Figure S6.**
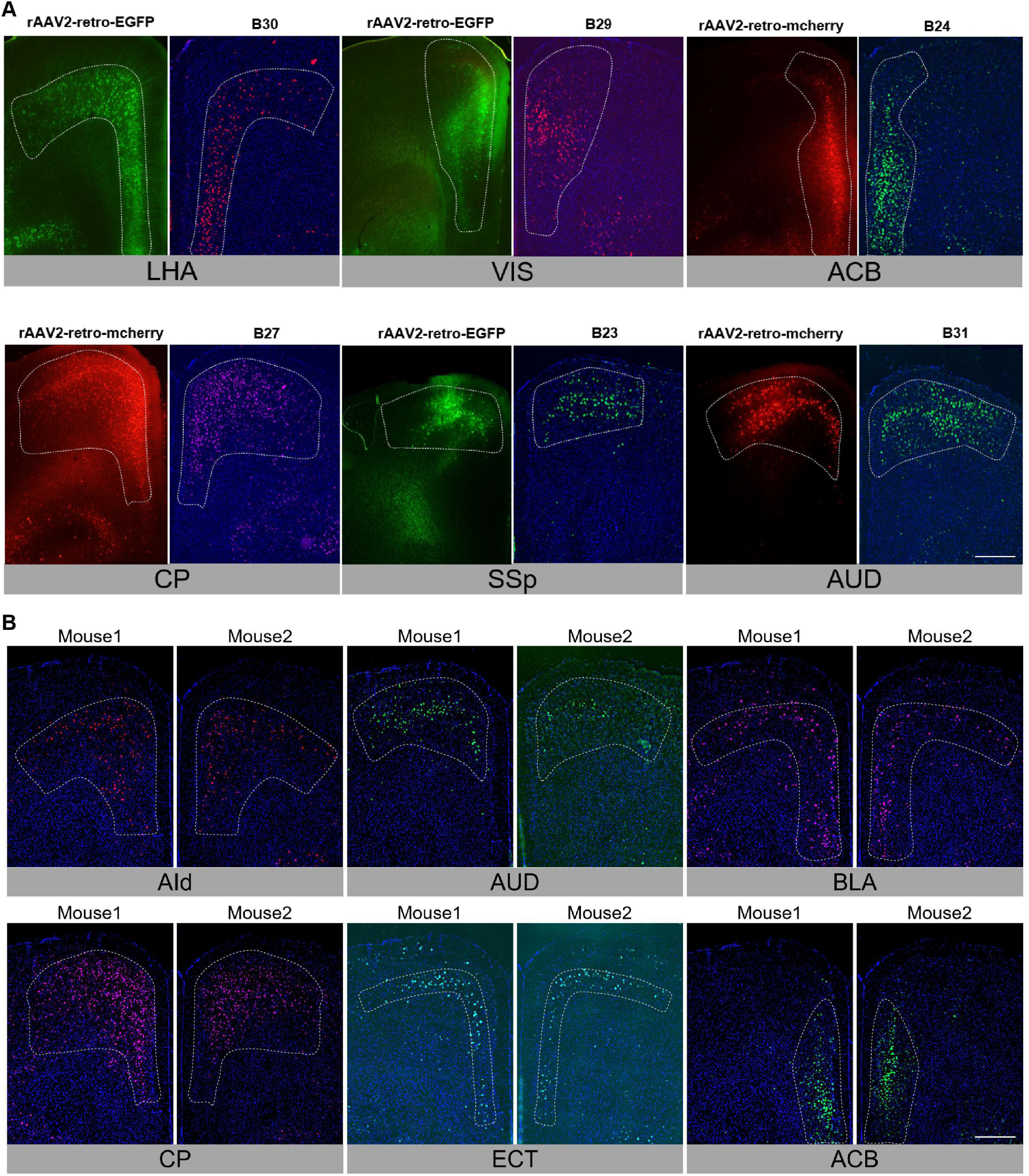
Comparison the barcode signal distribution obtained by in situ sequencing with fluorescent rAAV2-retro tracing and measurements in another mouse. **(A)** rAAV2-retro expressing mcherry or EGFP were injected into the target region (LHA, VIS, ACB, CP, SSp, AUD) and compare the distribution of fluorescent protein (right) with In situ sequencing signal of barcode (left). Scale bar: 2 mm. **(B)** Repeat measurements on six target regions (AId, AUD, BLA, CP, ECT, ACB) in another mouse using in-situ sequencing via SPIDER-seq. Scale bar: 2 mm.

**Figure S7.**
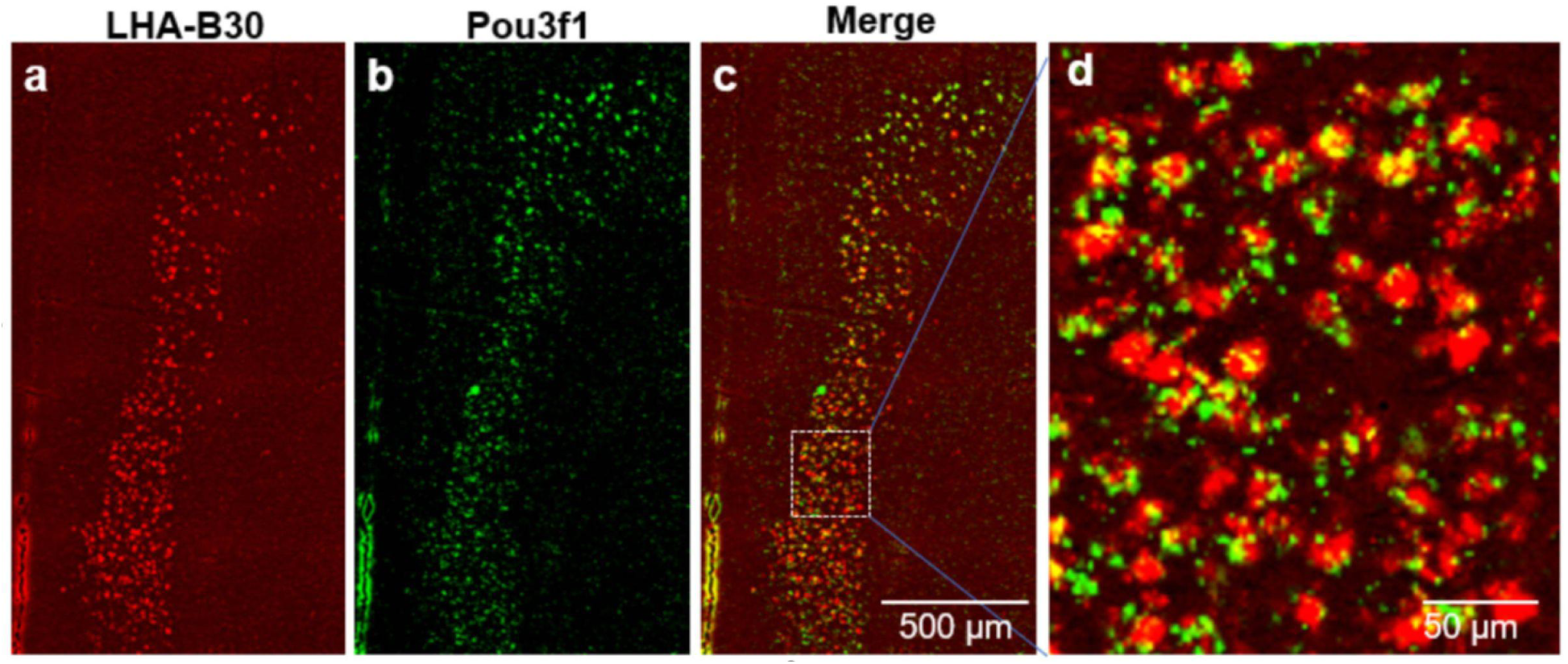
Barcode that retrogradely tracing from LHA is merged with the pyramidal tract neuron marker gene (*Pou3f1*) **(A)** In situ sequencing signal of LHA barcode. **(B)** In situ sequencing signal of L5 PT Marker gene: *Pou3f1*. **(C)** Merge of LHA barcode and *Pou3f1*. Scale bar: 500 µm. **(D)** Magnified view of the white boxed area in c. Scale bar: 50 µm.

**Figure S8.**
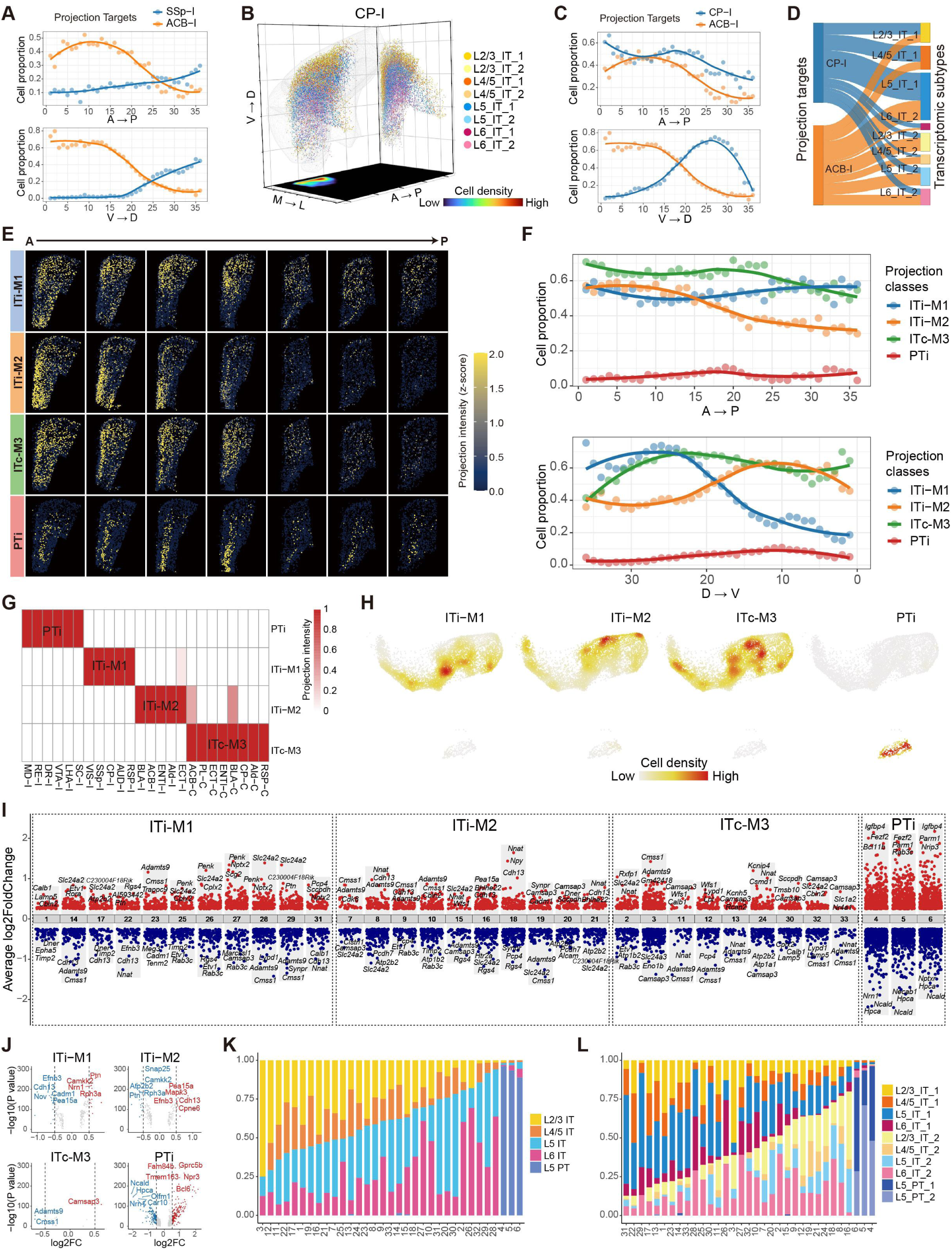
Deciphering the spatial and transcriptomic configuration of PFC projectome by SPIDER-Seq. **(A)** Distribution curves of PFC neurons projecting to ipsilateral ACB-I and SSp-I along the anterior-posterior axis (top) and the ventral-dorsal axis (bottom), respectively. **(B)** 3D visualization of the spatial distribution of PFC neurons projecting to ipsilateral CP-I, colored by transcriptomic subtypes. Right grid shows all neurons superimposed on the coronal plane. Bottom grid shows the density of all neurons on the transverse plane. **(C)** Distribution curves of PFC neurons projecting to ipsilateral CP-I and ACB-I along the anterior-posterior axis (top) and the ventral-dorsal axis (bottom), respectively. **(D)** Sankey diagram shows the transcriptomic subtypes composition of PFC neurons projecting to CP-I and ACB-I, respectively. **(E)** Spatial distribution of four projection classes along the anterior-posterior axis. **(F)** Spatial distribution curves of four projection classes along the anterior-posterior axis (top) and the ventral-dorsal axis (bottom). **(G)** Heatmap of the projection intensity of four projection classes to 24 targets. **(H)** Density scatter visualization on UMAP of four projection classes. The color scale represents the density of projection neurons. **(I)** Differentially expressed genes (DEGs) of 33 projection clusters. **(J)** Volcano plot showing DEGs in four PFC projection classes. **(K)** Transcriptome composition of 33 PFC projection clusters, grouped by transcriptomic cell layers. **(L)** Transcriptome composition of 33 PFC projection clusters, grouped by transcriptomic cell subtypes.

**Figure S9.**
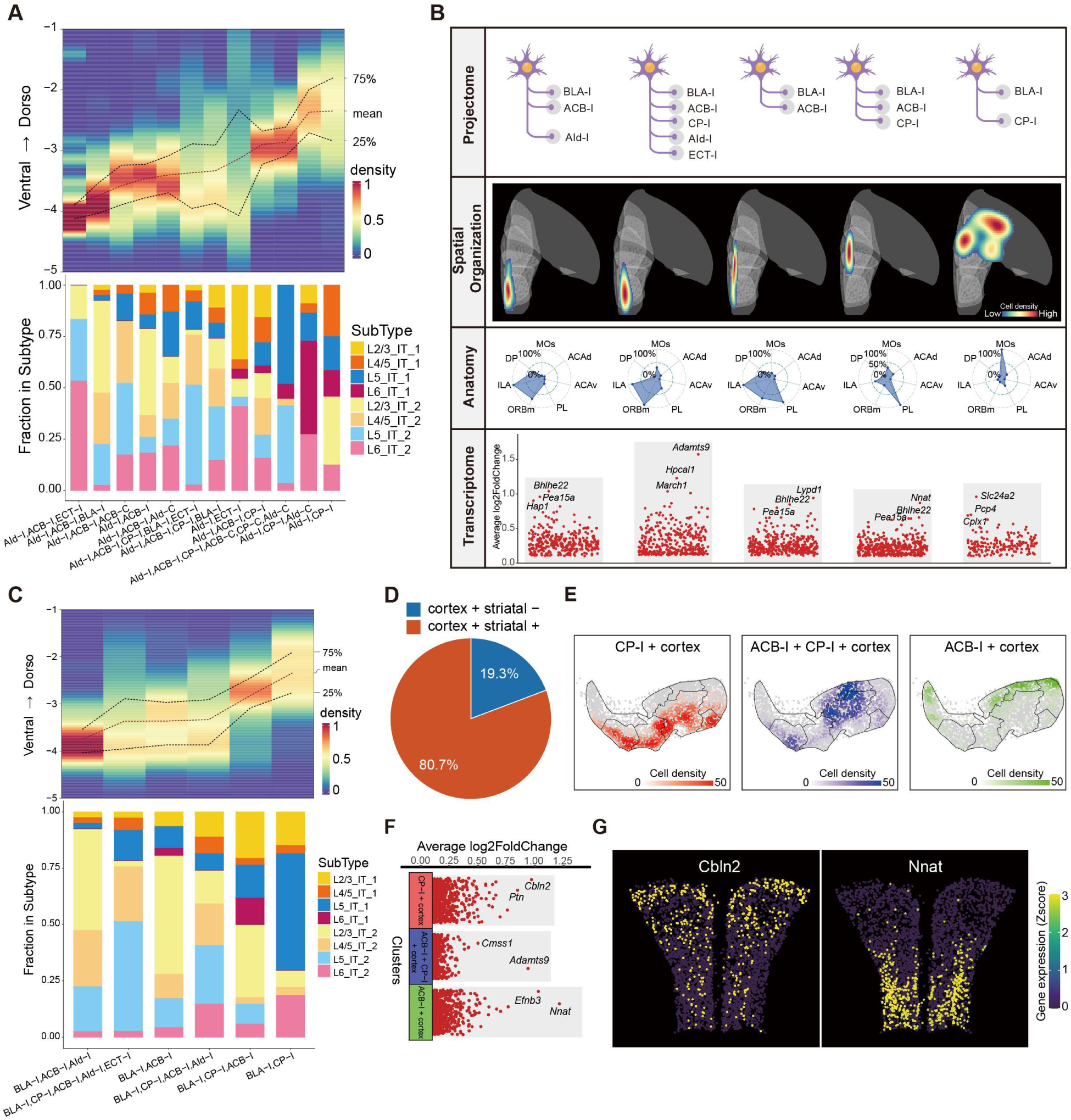
Organization pattern of PFC IT projection neurons. **(A)** Spatial distribution of neurons in different motif targeting AId-I along the ventral-dorsal axis (top), and transcriptome composition (bottom). **(B)** transcriptomics and spatial information of different PFC projection motifs projecting to ipsilateral BLA. Top, target nuclei. Middle, differentially expressed genes. Bottom, spatial distribution. **(C)** Spatial distribution of neurons in different motifs targeting BLA-I along the ventral-dorsal axis axis (top), and transcriptome composition (bottom). **(D)** The percentage of cortex+ striatal- and cortex+ striatal+ projection neurons. **(E)** Transcriptomic UMAP distribution of CP-I+Cortex (or BLA) (left), ACB-I+Cortex (or BLA) (middle), and ACB-I+CP-I+Cortex (or BLA) (right) projection motifs. **(F)** DEGs of CP-I+Cortex (or BLA) (top), ACB-I+Cortex (or BLA) (middle), and ACB-I+CP-I+Cortex (or BLA) (bottom) projection motifs. **(G)** Spatial expression of DEGs in (F) (Bregma: 2.1mm).

**Figure S10.**
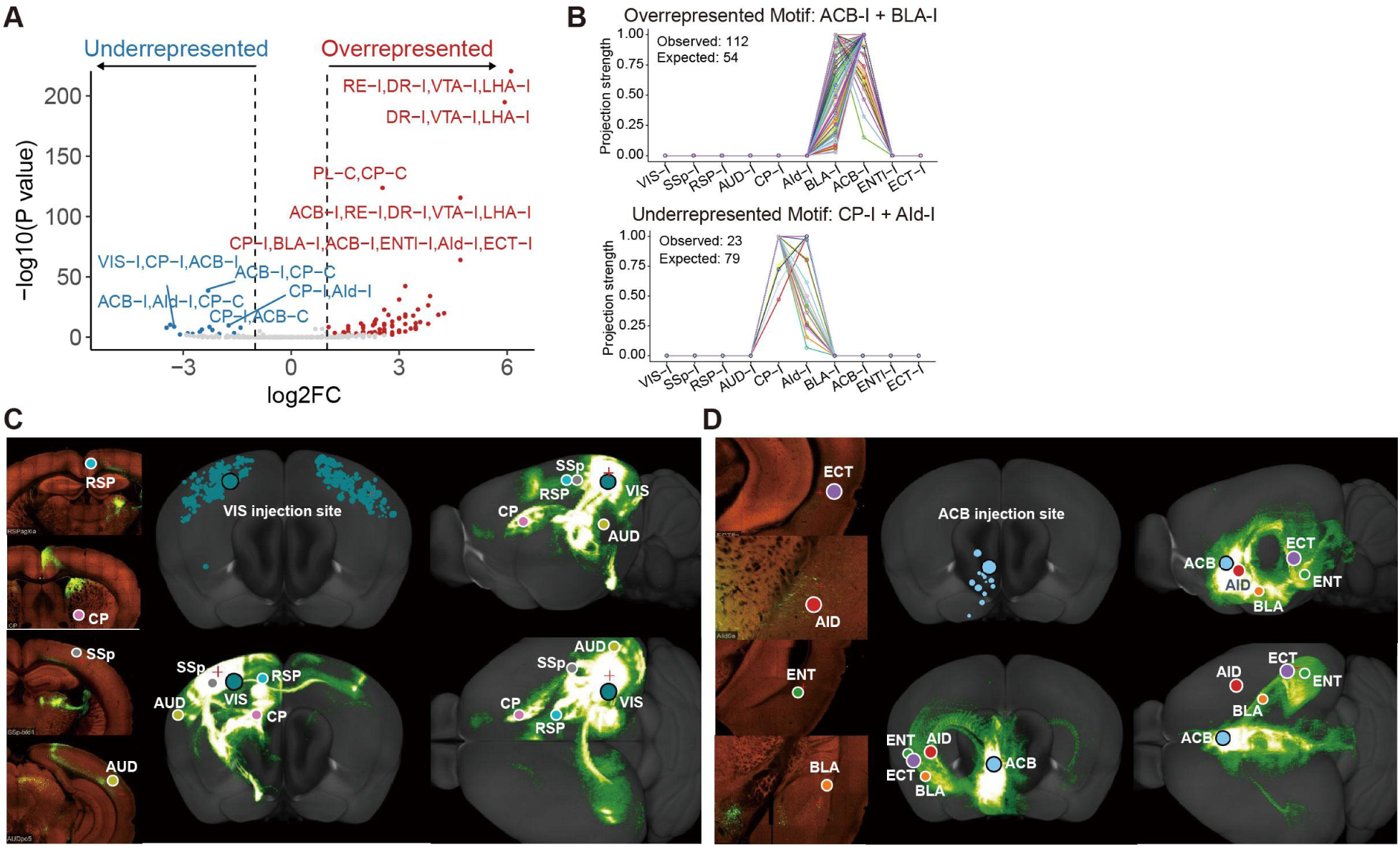
Circuit connections between PFC downstream nuclei. **(A)** The volcano plot shows the under-represented and over-represented projection motifs compared to null model. **(B)** Projection patterns of all individual neurons in the over-represented motif (ACB-I + BLA-I) and the under-represented motif (CP-I + AId-I). **(C)** Circuit connections between downstream targets of ITi-M1. Data from the Allen Mouse Brain Connectivity Atlas. **(D)** Circuit connections between downstream targets of ITi-M2. Data from the Allen Mouse Brain Connectivity Atlas.

**Figure S11.**
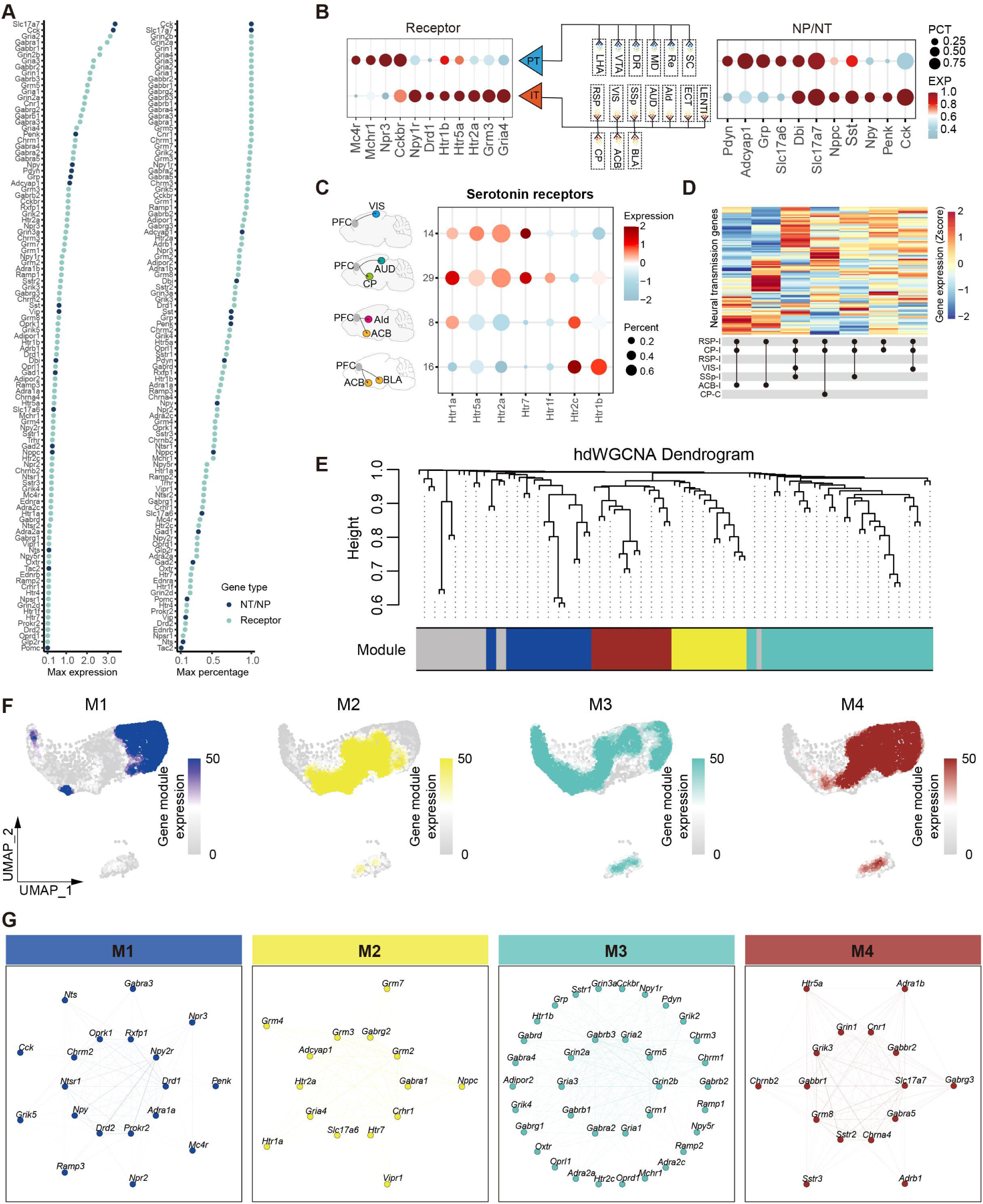
Expression characteristics of neural signaling molecule. **(A)** The maximum expression levels (left) and cell percentage (right) of neural signal molecules (neurotransmitter (NT)/neuropeptide (NP) and receptor genes) in different PFC projection clusters. **(B)** Differential neural signal transmission flow between IT and PT projection neurons of PFC. Dotplot showing the expression patterns of receptor genes (left) and neurotransmitter/neuropeptide genes (right) in IT/PT projection neurons. The middle panel showing the projection pattern of IT/PT projection neurons. **(C)** Different projection clusters expressing diverse serotonin receptor subtypes. **(D)** Different projection motifs target RSP-I have different neural signaling molecules expression patterns. **(E)** hdWGCNA dendrogram of the co-expression network of molecules related to neuronal signaling molecules using hdWGCNA. **(F)** UMAP colored by the module eigengenes (MEs) for the four gene modules. **(G)** Co-expression plots for the four gene modules.

**Figure S12.**
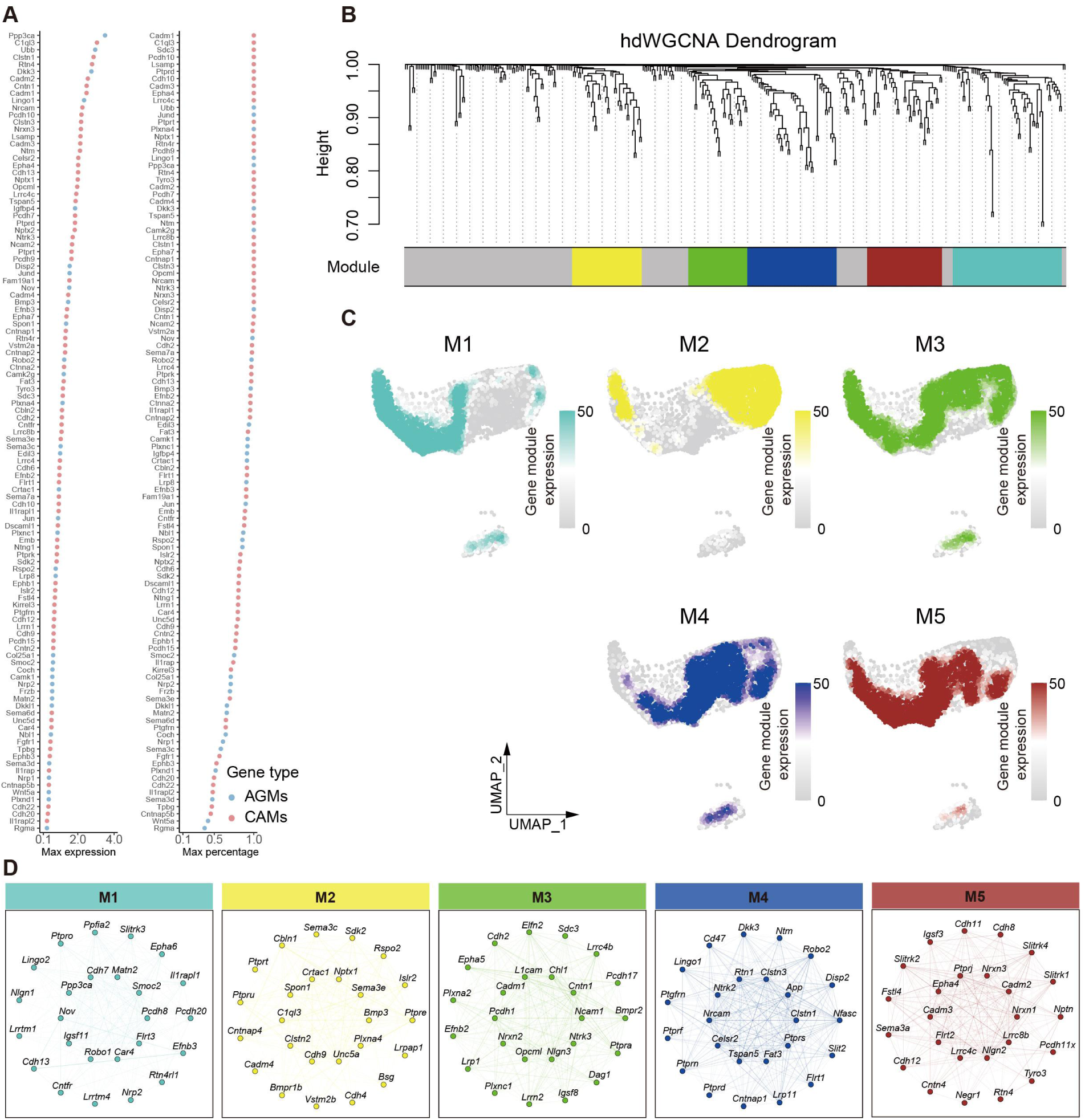
Expression characteristics of neuronal circuit wiring related genes. **(A)** The maximum expression levels (left) and cell percentage (right) of axon guidance molecules (AGMs) and cadherin molecules (CAMs) genes in different projection neurons. **(B)** hdWGCNA dendrogram of the co-expression network of molecules related to neuronal circuit wiring using hdWGCNA. **(C)** UMAP colored by the module eigengenes (MEs) for the five gene modules. **(D)** Co-expression plots for the five gene modules.

**Figure S13.**
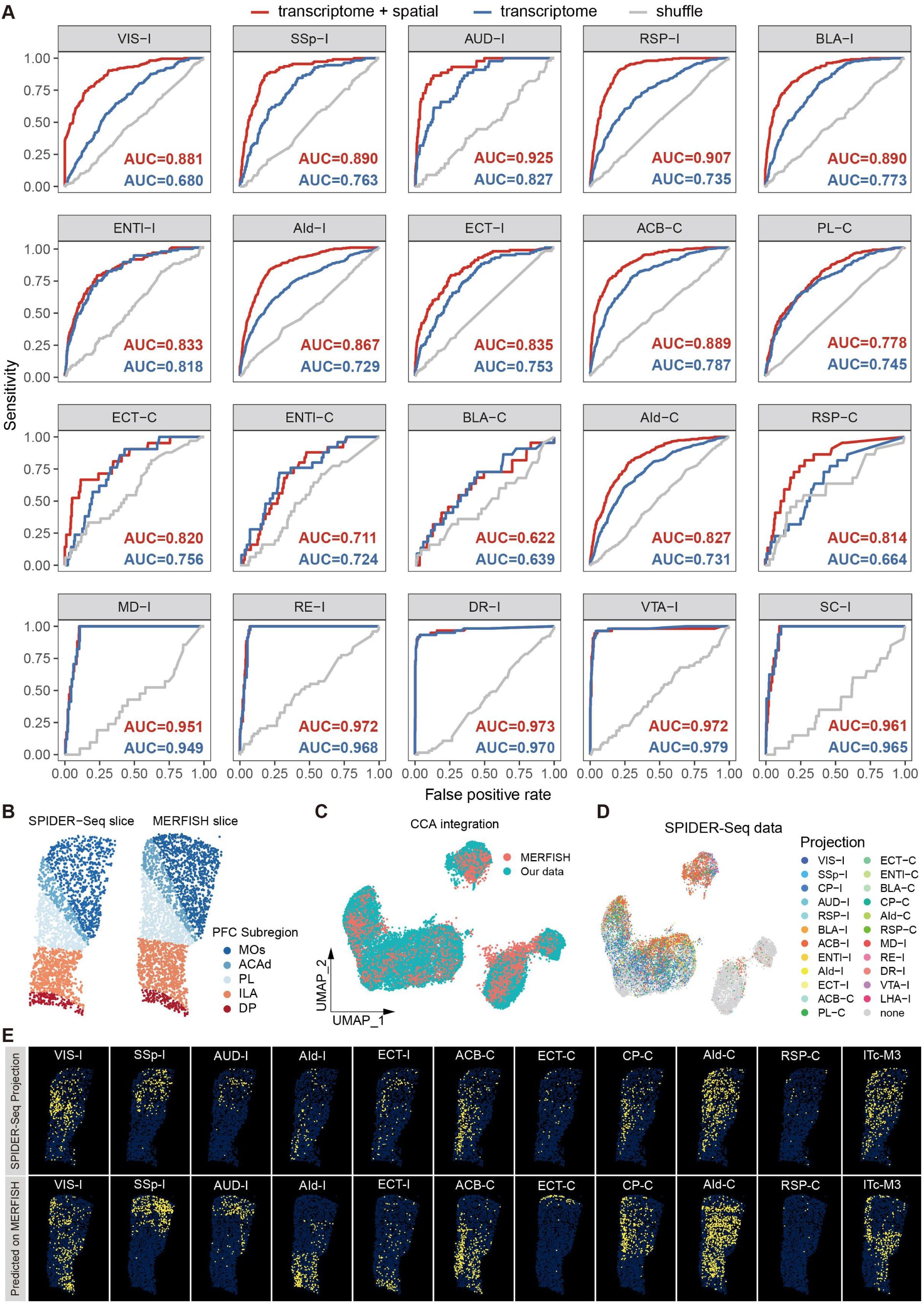
Prediction of neuron projection by integrated gene profile and spatial location information by machine learning. **(A)** ROC curves for predicted projection targets. The red curves utilize both transcriptome and spatial information as input, the blue curves utilize only transcriptome as input, and the gray curves are random shuffle control. **(B)** An example PFC slice from our SPIDER-Seq data (left) and the corresponding slice in MERFISH data (right), colored by PFC subregion. **(C)** UMAP visualization of our SPIDER-Seq data and MERFISH data after CCA integration. **(D)** UMAP visualization of our SPIDER-Seq data, with cells colored by the projection. **(E)** The spatial distribution of neuron to different projection targets measured by SPIDER-seq (top), and the putative distributions of neuron to different targets predicted by our machine learning model based on MERFISH data (bottom) (Bregma: 1.78mm).

**Figure S14.**
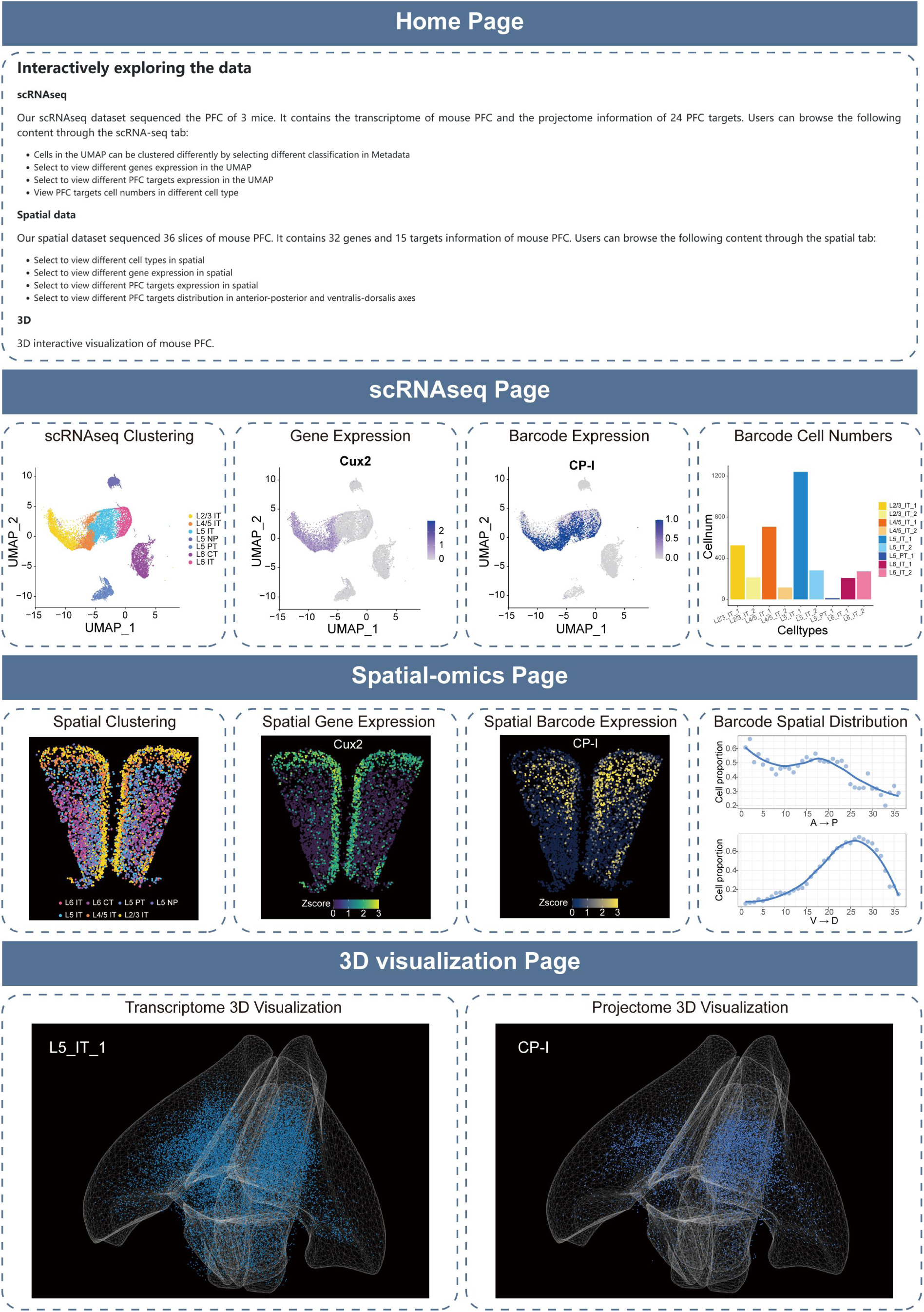
Overview of the SPIDER-web. SPIDER-web is a shiny application that allows users to interactively access our data. Home Page provides information about our project and how to use it interactively. Users can access our scRNAseq data through scRNAseq Page, and access our spatial-omics data through Spatial Page. 3D visualization Page provides interactive 3D visualization of the PFC transcriptome and projectome.

## REFERENCES

1. Seguin C, Sporns O, Zalesky A. Brain network communication: concepts, models and applications. Nature reviews Neuroscience. 2023; 24(9): 557–574. doi: 10.1038/s41583-023-00718-5

2. Suárez LE, Mihalik A, Milisav F et al. Connectome-based reservoir computing with the conn2res toolbox. Nature communications. 2024; 15(1): 656. doi: 10.1038/s41467-024-44900-4

3. Oh SW, Harris JA, Ng L et al. A mesoscale connectome of the mouse brain. Nature. 2014; 508(7495): 207-214. doi: 10.1038/nature13186

4. Kunst M, Laurell E, Mokayes N et al. A Cellular-Resolution Atlas of the Larval Zebrafish Brain. Neuron. 2019; 103(1): 21–38.e25. doi: 10.1016/j.neuron.2019.04.034

5. Scheffer LK, Xu CS, Januszewski M et al. A connectome and analysis of the adult Drosophila central brain. eLife. 2020; 9. doi: 10.7554/eLife.57443

6. Dorkenwald S, Matsliah A, Sterling AR et al. Neuronal wiring diagram of an adult brain. Nature. 2024; 634(8032): 124–138. doi: 10.1038/s41586-024-07558-y

7. Avila B, Augusto P, Hashemi A et al. Symmetries and synchronization from whole-neural activity in the Caenorhabditis elegans connectome: Integration of functional and structural networks. Proceedings of the National Academy of Sciences of the United States of America. 2025; 122(23): e2417850122. doi: 10.1073/pnas.2417850122

8. Zador AM, Dubnau J, Oyibo HK et al. Sequencing the connectome. PLoS biology. 2012; 10(10): e1001411. doi: 10.1371/journal.pbio.1001411

9. Luo L. Architectures of neuronal circuits. Science (New York, NY). 2021; 373(6559): eabg7285. doi: 10.1126/science.abg7285

10. Bienkowski MS, Bowman I, Song MY et al. Integration of gene expression and brain-wide connectivity reveals the multiscale organization of mouse hippocampal networks. Nature neuroscience. 2018; 21(11): 1628–1643. doi: 10.1038/s41593-018-0241-y

11. Yonehara K, Roska B. “MAPseq”-uencing Long-Range Neuronal Projections. Neuron. 2016; 91(5): 945–947. doi: 10.1016/j.neuron.2016.08.029

12. Han Y, Kebschull JM, Campbell RAA et al. The logic of single-cell projections from visual cortex. Nature. 2018; 556(7699): 51–56. doi: 10.1038/nature26159

13. Chen X, Sun YC, Zhan H et al. High-Throughput Mapping of Long-Range Neuronal Projection Using In Situ Sequencing. Cell. 2019; 179(3): 772–786.e719. doi: 10.1016/j.cell.2019.09.023

14. Huang L, Kebschull JM, Fürth D et al. BRICseq Bridges Brain-wide Interregional Connectivity to Neural Activity and Gene Expression in Single Animals. Cell. 2020; 182(1): 177–188.e127. doi: 10.1016/j.cell.2020.05.029

15. Qiu S, Hu Y, Huang Y et al. Whole-brain spatial organization of hippocampal single-neuron projectomes. Science (New York, NY). 2024; 383(6682): eadj9198. doi: 10.1126/science.adj9198

16. Sun YC, Chen X, Fischer S et al. Integrating barcoded neuroanatomy with spatial transcriptional profiling enables identification of gene correlates of projections. Nature neuroscience. 2021; 24(6): 873–885. doi: 10.1038/s41593-021-00842-4

17. Tasic B, Yao Z, Graybuck LT et al. Shared and distinct transcriptomic cell types across neocortical areas. Nature. 2018; 563(7729): 72–78. doi: 10.1038/s41586-018-0654-5

18. Cheung V, Chung P, Bjorni M et al. Virally encoded connectivity transgenic overlay RNA sequencing (VECTORseq) defines projection neurons involved in sensorimotor integration. Cell reports. 2021; 37(12): 110131. doi: 10.1016/j.celrep.2021.110131

19. Zhang Z, Zhou J, Tan P et al. Epigenomic diversity of cortical projection neurons in the mouse brain. Nature. 2021; 598(7879): 167–173. doi: 10.1038/s41586-021-03223-w

20. Zhou J, Zhang Z, Wu M et al. Brain-wide correspondence of neuronal epigenomics and distant projections. Nature. 2023; 624(7991): 355–365. doi: 10.1038/s41586-023-06823-w

21. Zhao Q, Yu CD, Wang R et al. A multidimensional coding architecture of the vagal interoceptive system. Nature. 2022; 603(7903): 878–884. doi: 10.1038/s41586-022-04515-5

22. Xu P, Peng J, Yuan T et al. High-throughput mapping of single-neuron projection and molecular features by retrograde barcoded labeling. eLife. 2024; 13. doi: 10.7554/eLife.85419

23. Anastasiades PG, Carter AG. Circuit organization of the rodent medial prefrontal cortex. Trends in neurosciences. 2021; 44(7): 550–563. doi: 10.1016/j.tins.2021.03.006

24. Le Merre P, Ährlund-Richter S, Carlén M. The mouse prefrontal cortex: Unity in diversity. Neuron. 2021; 109(12): 1925–1944. doi: 10.1016/j.neuron.2021.03.035

25. Negrón-Oyarzo I, Aboitiz F, Fuentealba P. Impaired Functional Connectivity in the Prefrontal Cortex: A Mechanism for Chronic Stress-Induced Neuropsychiatric Disorders. Neural plasticity. 2016; 2016: 7539065. doi: 10.1155/2016/7539065

26. Friedman NP, Robbins TW. The role of prefrontal cortex in cognitive control and executive function. Neuropsychopharmacology : official publication of the American College of Neuropsychopharmacology. 2022; 47(1): 72–89. doi: 10.1038/s41386-021-01132-0

27. Bhattacherjee A, Djekidel MN, Chen R et al. Cell type-specific transcriptional programs in mouse prefrontal cortex during adolescence and addiction. Nature communications. 2019; 10(1): 4169. doi: 10.1038/s41467-019-12054-3

28. Ma S, Skarica M, Li Q et al. Molecular and cellular evolution of the primate dorsolateral prefrontal cortex. Science (New York, NY). 2022; 377(6614): eabo7257. doi: 10.1126/science.abo7257

29. Bhattacherjee A, Zhang C, Watson BR et al. Spatial transcriptomics reveals the distinct organization of mouse prefrontal cortex and neuronal subtypes regulating chronic pain. Nature neuroscience. 2023; 26(11): 1880–1893. doi: 10.1038/s41593-023-01455-9

30. Lui JH, Nguyen ND, Grutzner SM et al. Differential encoding in prefrontal cortex projection neuron classes across cognitive tasks. Cell. 2021; 184(2): 489–506.e426. doi: 10.1016/j.cell.2020.11.046

31. Gao L, Liu S, Gou L et al. Single-neuron projectome of mouse prefrontal cortex. Nature neuroscience. 2022; 25(4): 515–529. doi: 10.1038/s41593-022-01041-5

32. Wu X, Xu W, Deng L et al. Spatial multi-omics at subcellular resolution via high-throughput in situ pairwise sequencing. Nature biomedical engineering. 2024; 8(7): 872–889. doi: 10.1038/s41551-024-01205-7

33. Biancalani T, Scalia G, Buffoni L et al. Deep learning and alignment of spatially resolved single-cell transcriptomes with Tangram. Nature methods. 2021; 18(11): 1352–1362. doi: 10.1038/s41592-021-01264-7

34. de Kloet SF, Bruinsma B, Terra H et al. Bi-directional regulation of cognitive control by distinct prefrontal cortical output neurons to thalamus and striatum. Nature communications. 2021; 12(1): 1994. doi: 10.1038/s41467-021-22260-7

35. Deng B, Li Q, Liu X et al. Chemoconnectomics: Mapping Chemical Transmission in Drosophila. Neuron. 2019; 101(5): 876–893.e874. doi: 10.1016/j.neuron.2019.01.045

36. Ripoll-Sánchez L, Watteyne J, Sun H et al. The neuropeptidergic connectome of C. elegans. Neuron. 2023; 111(22): 3570–3589.e3575. doi: 10.1016/j.neuron.2023.09.043

37. Wallace ML, Sabatini BL. Synaptic and circuit functions of multitransmitter neurons in the mammalian brain. Neuron. 2023; 111(19): 2969–2983. doi: 10.1016/j.neuron.2023.06.003

38. Zhao W, Johnston KG, Ren H et al. Inferring neuron-neuron communications from single-cell transcriptomics through NeuronChat. Nature communications. 2023; 14(1): 1128. doi: 10.1038/s41467-023-36800-w

39. Granger AJ, Wallace ML, Sabatini BL. Multi-transmitter neurons in the mammalian central nervous system. Current opinion in neurobiology. 2017; 45: 85–91. doi: 10.1016/j.conb.2017.04.007

40. Chen MB, Jiang X, Quake SR et al. Persistent transcriptional programmes are associated with remote memory. Nature. 2020; 587(7834): 437–442. doi: 10.1038/s41586-020-2905-5

41. Dutta S, Beaver J, Halcomb CJ et al. Dissociable roles of the nucleus accumbens core and shell subregions in the expression and extinction of conditioned fear. Neurobiology of stress. 2021; 15: 100365. doi: 10.1016/j.ynstr.2021.100365

42. Sun W, Liu Z, Jiang X et al. Spatial transcriptomics reveal neuron-astrocyte synergy in long-term memory. Nature. 2024; 627(8003): 374–381. doi: 10.1038/s41586-023-07011-6

43. Ortiz C, Navarro JF, Jurek A et al. Molecular atlas of the adult mouse brain. Science advances. 2020; 6(26): eabb3446. doi: 10.1126/sciadv.abb3446

44. Gergues MM, Han KJ, Choi HS et al. Circuit and molecular architecture of a ventral hippocampal network. Nature neuroscience. 2020; 23(11): 1444–1452. doi: 10.1038/s41593-020-0705-8

45. Sun L, Tang Y, Yan K et al. Differences in neurotropism and neurotoxicity among retrograde viral tracers. Molecular neurodegeneration. 2019; 14(1): 8. doi: 10.1186/s13024-019-0308-6

46. Saunders A, Macosko EZ, Wysoker A et al. Molecular Diversity and Specializations among the Cells of the Adult Mouse Brain. Cell. 2018; 174(4): 1015–1030.e1016. doi: 10.1016/j.cell.2018.07.028

47. Hao Y, Hao S, Andersen-Nissen E et al. Integrated analysis of multimodal single-cell data. Cell. 2021; 184(13): 3573–3587.e3529. doi: 10.1016/j.cell.2021.04.048

48. McGinnis CS, Murrow LM, Gartner ZJ. DoubletFinder: Doublet Detection in Single-Cell RNA Sequencing Data Using Artificial Nearest Neighbors. Cell systems. 2019; 8(4): 329–337.e324. doi: 10.1016/j.cels.2019.03.003

49. Danecek P, Bonfield JK, Liddle J et al. Twelve years of SAMtools and BCFtools. GigaScience. 2021; 10(2). doi: 10.1093/gigascience/giab008

50. Stringer C, Wang T, Michaelos M et al. Cellpose: a generalist algorithm for cellular segmentation. Nature methods. 2021; 18(1): 100–106. doi: 10.1038/s41592-020-01018-x

51. Fürth D, Vaissière T, Tzortzi O et al. An interactive framework for whole-brain maps at cellular resolution. Nature neuroscience. 2018; 21(1): 139–149. doi: 10.1038/s41593-017-0027-7

52. Morabito S, Reese F, Rahimzadeh N et al. hdWGCNA identifies co-expression networks in high-dimensional transcriptomics data. Cell reports methods. 2023; 3(6): 100498. doi: 10.1016/j.crmeth.2023.100498

